# A system to decipher the origins and relationships of all arthropod outgrowths

**DOI:** 10.1101/2021.01.20.427514

**Authors:** Heather S. Bruce, Nipam H. Patel

**Affiliations:** University of British Columbia, Vancouver, BC V6T1Z4 Canada; Marine Biological Laboratory, 7 MBL St. Woods Hole, MA 02543 USA; University of Chicago, Organismal Biology & Anatomy, 1027 E 57th Street, Chicago, IL 60637 USA

**Keywords:** Arthropod, appendage, homology, *Parhyale*, *Tribolium*, *Acanthoscurria*

## Abstract

Arthropods (insects, “crustaceans”, myriapods, and chelicerates) display a fascinating diversity of ectodermal structures like plates, horns, helmets, knobs, carapaces, mimicry outgrowths, and wings. The origins and relationships of these structures has tantalized researchers for over a century: did these structures arise de novo in each lineage, or do they emerge from shared, ancestral primordia present in some or even all arthropods? One way to begin answering this is to assess the position and context of these structures on arthropod bodies: do these structures emerge from proximal embryonic leg segments that were converted into the body wall, or do they represent dorsal, non-leg-derived structures? Here, the expression of *pannier*, *araucan*, and *drumstick* – genes previously shown to distinguish proximal leg segments in crustaceans and insects – are examined in a chelicerate representative, the tarantula *Acanthoscurria*. This gene expression comparison, together with over a century of gene functional, morphological, embryological, and paleontological data, suggests that all arthropod leg segments correspond to each other in a one-to-one fashion, but that in many arthropod lineages, the base of the ancestral leg in the embryo flattens and expands to form the body wall of the adult. This would mean that many arthropod outgrowths which appear to stand on the lateral body wall, such as wings, tergal plates, and gin traps, are derived from the ancestral/embryonic leg base, and likely arose from shared, ancestral primordia present on this leg base in all arthropods. The analysis detailed here means that a simple three- or four-gene in situ expression experiment with *pannier*, *araucan*, and *drumstick* can elucidate the homologies of any arthropod ectodermal structure of interest, including beetle horns, treehopper helmets, and more.

## Introduction

The diversity of arthropod appendages has facilitated their success across every biome. There are four main groups of living arthropods: chelicerates (spiders, etc.), myriapods (millipedes, etc.), “crustaceans” (i.e. the paraphyletic clade of non-insect pancrustaceans, such as shrimps, etc.), and insects (beetles, etc.). Each group has different numbers of and names for their anatomical leg segments (Fig. 1)^1–5^. Understanding how the leg segments of the four different arthropod groups correspond to each other has tantalized researchers for over a century^2,4,6–11^, because this would answer generations of speculation about the origins and possible relationships of fascinating arthropod structures like wings, gills, helmets, and horns^6,12–23^.

**Fig. 1.**
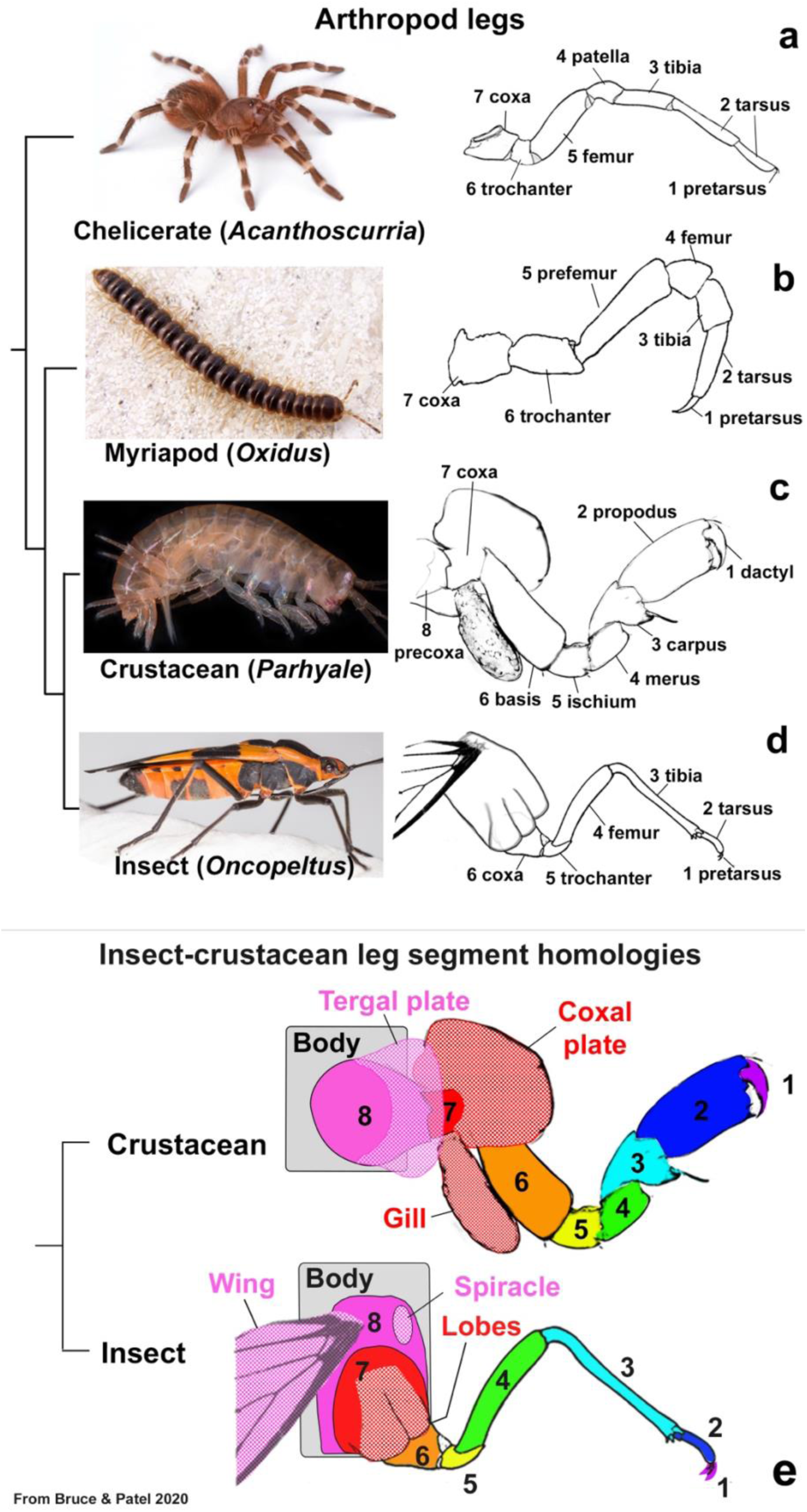
Arthropod legs. **a-d.** Chelicerates, myriapods, crustaceans, and insects have different numbers, shapes, and names for their leg segments. **e.** Leg segment correspondence between insect and crustacean leg segments. Colors indicate proposed homologies. Exites indicated with checker pattern. Panel modified from Bruce and Patel^24^. Phylogeny based on Lozano-Fernandez 2019^111^ and Bernot 2023^112^. *Acanthoscurria* and *Oxidus* images from Wikipedia. Leg drawings adapted from Snodgrass 1952^1^, except *Parhyale* leg, original image.

In our 2020 paper, we sought to understand the leg segment relationships between two of the four groups of arthropods, insects and crustaceans^24^. To do so, we compared a century of previous morphological and embryological work with the expression and function of several leg-and wing-patterning genes between insects and crustaceans, using original and published data.

Authors have historically proposed a variety of leg segment deletions, duplications, and/or fusions to account for the different numbers of leg segments in the four arthropod groups^2,4,8,14,25–32^. However, our work (Fig. 1e) suggested that insect and crustacean leg segments have a simple, one-to-one correspondence, and the apparent loss of two leg segments in insects is due to the embryonic expansion and flattening of two proximal leg segments to form the adult insect lateral body wall (pleura and lateral tergum; also called subcoxa and precoxa/epicoxa, respectively)^14,33–37^. We found that two genes – the Iroquois complex gene *araucan* (*ara*) and the GATA factor *pannier* (*pnr*) – distinguish these proximal leg segments even if, as in insects, the leg segments now function as body wall^24,38–45^. This molecular comparison supported previous morphological, embryological, and fossil studies which suggested that arthropods ancestrally have a total of 8 leg segments^2,4,8,34,46,47^.

We therefore wondered whether this model held true for all arthropods. There is currently no gene function data for myriapods, so we focused on chelicerates. To determine whether chelicerates with 7 leg segments had converted an ancestral/embryonic proximal leg segment into the body wall, we investigated the expression of *pnr*, *ara,* and the arthropod joint marker *drumstick* in the tarantula spider *Acanthoscurria geniculata*. We then placed our results within the broader context of over a century of morphological, embryological, fossil, and molecular studies in all arthropods.

## Results

Extinct arthropods at the base of the arthropod phylogenetic tree, as well as early-branching chelicerates^48,49^ like sea spiders, camel spiders, and some mites, have 8 true (eudesmatic) leg segments^4,46^, but later-branching chelicerates like spiders and harvestman have only 7 apparent leg segments^4^. This begs the question of whether and how the ancestral 8^th^ leg segment was lost in these chelicerates: was this ancestral leg segment deleted, or was it converted into the body wall as insects have done?^8,24,28,33,34,36,47,50–54^

To elucidate the composition of proximal leg segments in chelicerates, the expression of *pnr*, *ara,* and *Dll* was compared between the crustacean *Parhyale*, the insect *Tribolium*, and the tarantula *Acanthoscurria geniculata* (Fig. 2)^24^. In *Drosophila*, *pannier* (*pnr*) patterns the dorsal midline, extra-embryonic membranes, and heart. Loss of *pnr* results in failure of embryonic dorsal closure, deletion of the dorsal midline ectoderm^43,55^, and failure of the dorsal mesoderm to become the heart^56^, whereas cells in the wing induced to express *pnr* form dorsal medial body wall (notum)^39^. Iroquois complex genes, including *araucan-caupolican* (*ara-caup*), pattern the left and right lateral notum in *Drosophila*, and loss transforms the lateral body wall (notum) into wing hinge^38,40,41,57^.

**Fig. 2.**
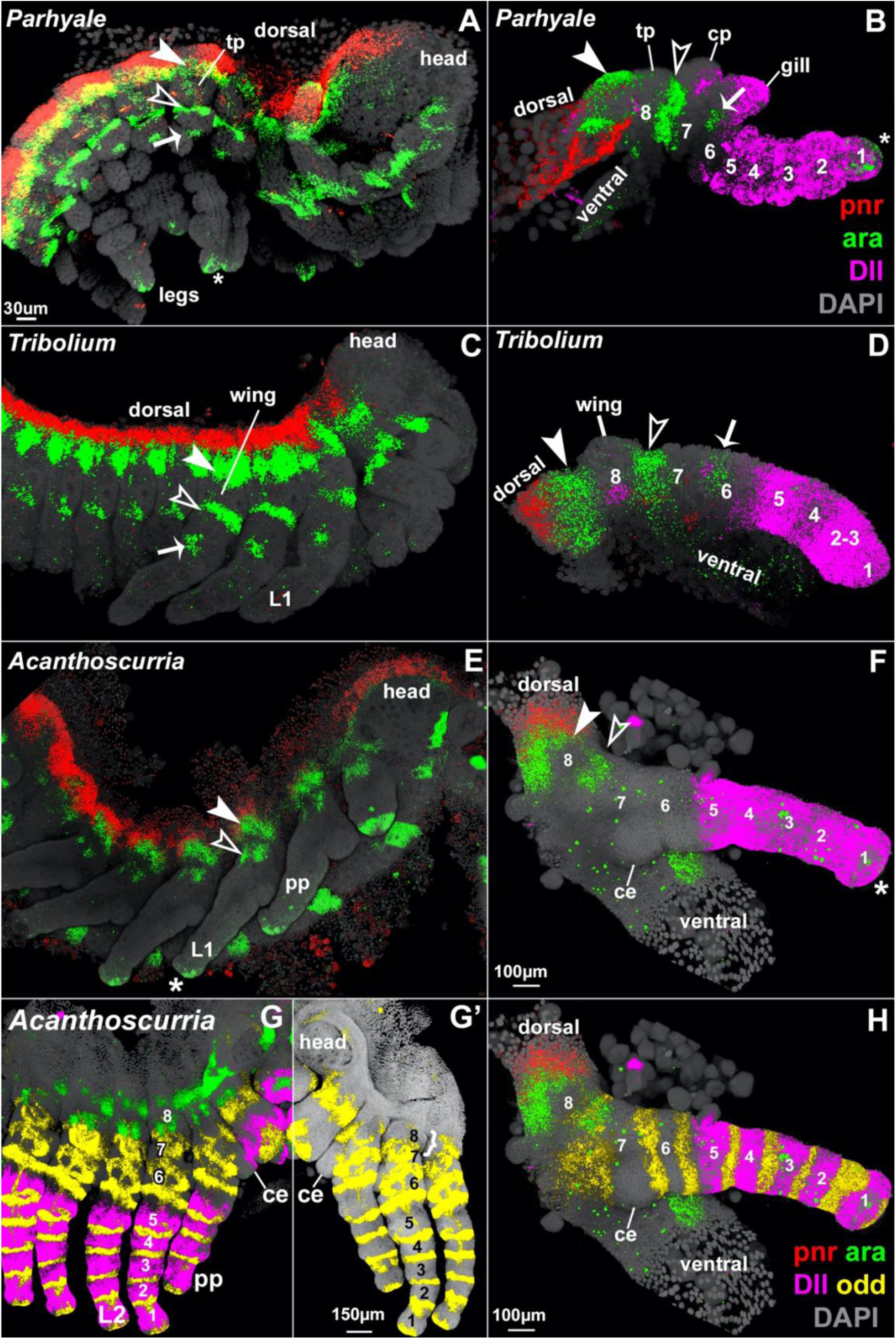
Elucidating proximal leg segments in arthropods. Dissected right half of *Parhyale* (A), *Tribolium* (C), and *Acanthoscurria* (E) embryos. Dissected leg of *Parhyale* (B), *Tribolium* (D), and *Acanthoscurria* (F) embryos. *pnr* (red), *ara* (green), *Dll* (pink), DAPI (grey). In all arthropods for which there is data, leg segments 1 through 5 are identified by *Dll* expression at the embryonic stage where leg joints first become visible^13,24,71,73,76,78,80,81,87–89,92,143,144,146,168–186^. *Dll* also expressed in *Parhyale* gill, coxal plate (cp) and tergal plate (tp). In all three arthropods, *pnr* marks the most dorsal domain. In all three arthropods, the two *ara* armband domains bracket a region proximal to leg segment 7. *ara* is also expressed in a smattering of ventral non-leg cells, and in *Parhyale* and *Acanthoscurria*, in the distal tip of the leg (*). *Tribolium* larvae have a fused tibia and tarsus, the tibiotarsus, here labelled 2-3^53^. In *Acanthoscurria*, leg segment 7 is easily identified by the coxal endite (ce) that bulges medially. Large dorsal cells in *Parhyale* and *Acanthoscurria* are yolk or extra-embryonic cells that exist prior to dorsal closure. (G – H) *Acanthoscurria drm* (yellow) is expressed in the distal region of each leg segment where the joint will later form. (G) Stage 12.5 *Acanthoscurria* embryos dissected away from yolk mass. Leg segment 7 is readily identified by the coxal endite. At this stage, proximal to leg segment 7, there is a leg-segment-like bulge (white curly brace), which expresses *drm* in its distal region (most visible in G’). By St 12.5, *pnr* expression in the walking legs and head is reduced and not visible through the other colors (compare with St 11.5 embryo in G). (H) dissected walking leg 1 from Stage 11.5 embryo where morphological bulges and subdivisions of the leg segments have not yet begun. *drm* encircles the distal region of each leg segment, including the hypothesized proximal 8^th^ leg segment. *Dll* and *pnr* in F, H in same channel but have non-overlapping domains, *Dll* false colored. See Fig. S2 for separate *Dll* and *pnr Acanthoscurria* embryos. A, C from Bruce and Patel 2020^24^

From the *Acanthoscurria* transcriptome^58^, OrthoFinder^59^ identified two paralogs of *pnr* in *Acanthoscurria* that were the reciprocal best blast hits to the single *pnr* ortholog in *Drosophila*, *Tribolium*, and *Parhyale* (Fig. S1). However, only one of these was expressed in *Acanthoscurria* embryos at the stages examined and was therefore presumed to be the conserved paralog of *pnr*. Additionally, OrthoFinder^59^ identified several Iroquois genes, but only one best *Acanthoscurria araucan* gene, and one best *Acanthoscurria Distal-less* (*Dll*) which were the closest to their respective orthologs in *Drosophila*, *Tribolium*, and *Parhyale*.

In order to facilitate the following comparisons, the names of each leg segment in insects, crustaceans, and chelicerates have been replaced with numbers 1 – 8, counting from the terminal claw (i.e. pretarsus, dactyl, etc.), as in Bruce and Patel 2020^24^, but the reader may refer to Fig. 1 for quick look-up. For example, the trochanter of chelicerates, the basis of crustaceans, and the coxa of insects are leg segment 6. For the insect representative used here, *Tribolium* (with has 6 moveable leg segments), the boundaries of the embryonic leg segments 7 and 8 that form the adult lateral body wall (pleura and lateral tergum, respectively) are based on molecular and morphological analyses in Kobayashi 2022^37^, as well as on the extensive literature regarding the “insect subcoxa theory”, such as the location of the paracoxal suture in Matsuda 1970^36^, Kobayashi 2013^34^, 2017^35^, and 2018^60^.

In *Parhyale*, *Tribolium,* and *Acanthoscurria, Dll* was found to be expressed in leg segments 1 – 5 and *pnr* expressed in the most dorsal tissue (Fig. 2). Thus, in all arthropods for which there is data, including the insects *Drosophila*^39,43,55,61,62^*, Tribolium*^24^, and *Bicyclus*^63^; the crustaceans *Parhyale*^24^*, Triops* (unpublished), and *Daphnia*^44^; the millipede *Glomeris*^64^; the chelicerate *Parasteatoda* (called GATA1)^65^; and here the chelicerate *Acanthoscurria*, *pnr* marks the “true” body wall (medial tergum) that is not derived from the leg base. In all arthropods examined to date, *ara* is expressed in four possible domains: a dorsal armband on proximal leg segment 8 that is adjacent to the *pnr* domain; a second armband at the joint between leg segments 7 and 8; a circular patch of expression on the lateral side of leg segment 6; and in the distal tip of leg segment 1. The crustaceans *Parhyale* and *Daphnia* express all four domains of *ara*^24,44^. In *Tribolium*, *ara* is expressed in the first three of these domains but not in the distal tip of the leg. In *Acanthoscurria*, *ara* is expressed in three of these domains: the dorsal armband adjacent to the *pnr* domain; the second armband on proximal leg segment 7; and in the distal tip of the leg. The lack of the lateral dot of *ara* in *Acanthoscurria* legs may be connected to the evolution of exopods within the pancrustaceans. Thus, as predicted by the leg segment alignment model in Bruce and Patel 2020 for insect and crustacean legs^24^, the two armbands of *ara* expression in *Acanthoscurria* bracket a region proximal to leg segment 7 (coxa) and adjacent to *pnr*. This suggests that *Acanthoscurria,* like *Parhyale*, *Daphnia*, and *Tribolium*, did not delete the ancestral, proximal 8^th^ leg segment, but rather converted it into the body wall.

To further test this hypothesis, the expression of *drumstick* was examined in *Acanthoscurria*. *Drumstick* is in the *odd-skipped* family of genes, which are expressed in the distal edge of each leg segment in all arthropods examined, including in *Parhyale*, *Tribolium*, and the chelicerate spider *Parastatoda*^63,66–69^. In particular, *drumstick* is expressed in a sharp line and only in the true (eudesmatic) leg joints, and not in the leg subdivisions that provide flexion such as those of the tarsus. Thus, expression of *drumstick* can be used to count the number of true leg segments present, and to molecularly distinguish true leg segments from superficial subdivisions. In *Drosophila*, all members of the *odd-skipped* family of genes – *odd-skipped* (*odd*), *brother of odd with entrails limited (bowl), sister of odd and bowl (sob), and drumstick (drm) –* have been shown to have similar overlapping expression patterns in the leg joints and to variably induce cells to buckle and form a flexible joint^70^. OrthoFinder^59^ identified a gene in *Acanthoscurria* and in *Parhyale* which were each the reciprocal best BLAST hit to *drm* in *Drosophila* and *Tribolium*. This *Acanthoscurria drumstick* was the reciprocal best BLAST hit to *Cupiennius* (spider) *odd-related* 3 (*odd-r3*)^66^. As in other arthropods, *Parhyale drm* is expressed in 7 stripes, one on the distal side of each of the 8 leg segments, denoting the future location of leg joints (Fig. S4)^67,68^. Similarly, in *Acanthoscurria, drm* is expressed in the expected distal regions of leg segments 1 – 7, but also in an additional ring proximal to leg segment 7 (Fig. 2G, H). This additional ring of *drm* notably occurs on the distal side of a leg-segment-like bulge, as expected if this bulge represents an embryonic 8^th^ leg segment. This leg-segment-like bulge is visible in other studies and other spider species, including *Acanthoscurria* embryos in Pechmann 2009^71^ Figure 3 and Pechmann 2020^72^ Figure 6; as well as in *Cupiennius* in Pechmann 2010 Figure 16, and Wolff 2011 Figures 11 and 12. Given that *drm* specifically marks the distal side of leg segments^70^, the ring of *drm* expression on the distal side of the leg-segment-like bulge suggests that it is indeed a leg segment. Together, the conserved expression of *pnr*, *ara,* and *drm* surrounding the embryonic leg-segment-like bulge suggest that *Acanthoscurria* has an additional, cryptic proximal 8^th^ leg segment.

**Fig. 3.**
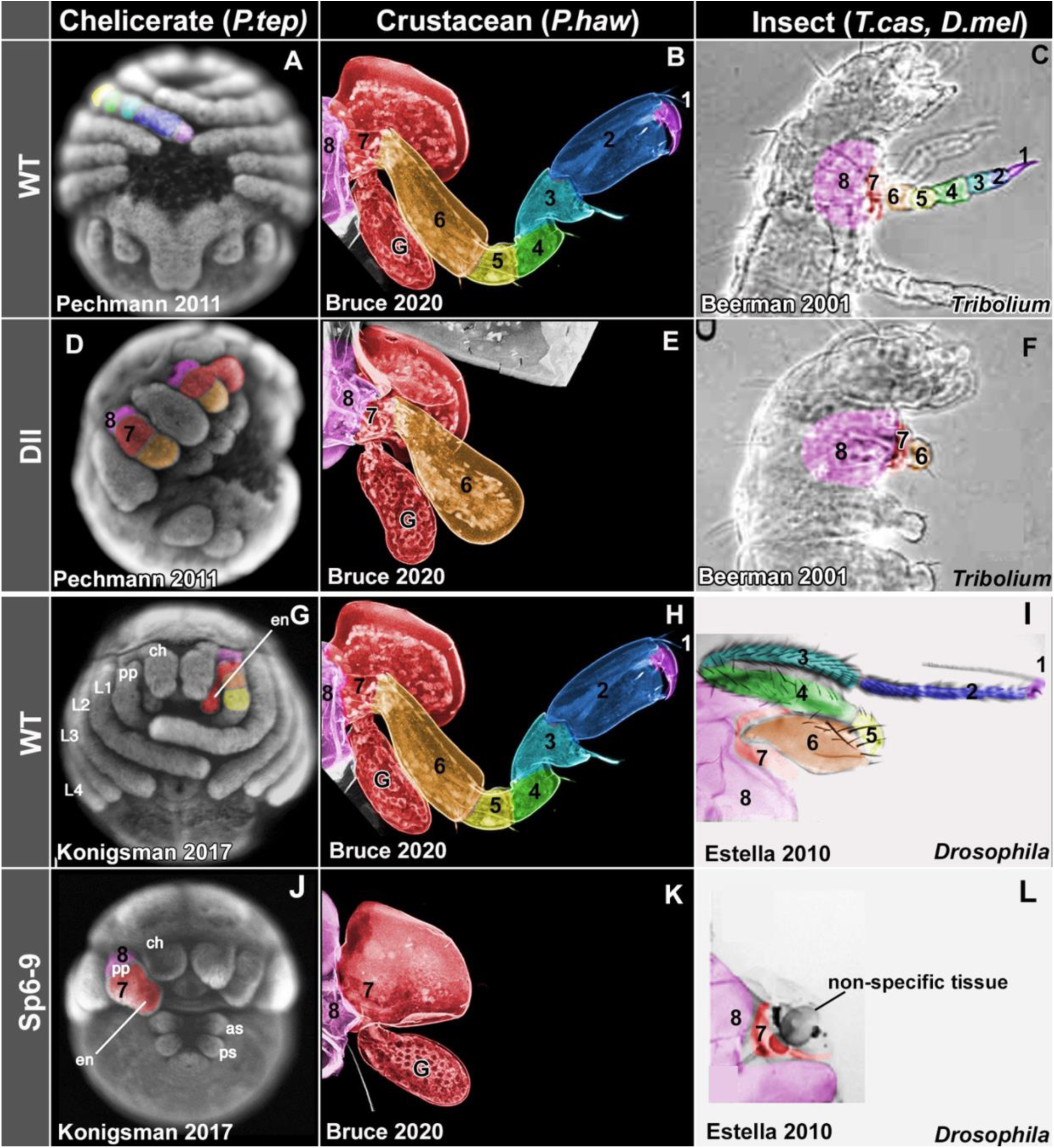
*Dll* and *Sp6-9* function in arthropods. In all arthropods examined, including the chelicerate *Parasteatoda*, crustacean *Parhyale*, and insect *Tribolium*, *Dll* is expressed in and required for the development of leg segments 1 - 5, counting from the distal end of the leg^24,79,85–90^. In all arthropods examined, including *Parasteatoda*, *Parhyale,* and *Drosophila*, *Sp6-9* is expressed in and required for the development of leg segments 1 – 6^24,77,82,91–93^. In *Parasteatoda* embryos, leg segment boundaries are best approximations based on embryo morphology; the respective author’s interpretations of their loss of function phenotypes, which were confirmed visually from their figures; as well as published gene expression boundaries for *Dll* and *Sp6-9* in *Parasteatoda* and other spiders and their respective author’s interpretations^71,73,79,80,84^. In C, F, boundaries of leg segments 7 and 8 that form the lateral body wall of *Tribolium* are based on molecular and morphological analyses in Kobayashi 2022^37^; the remnant of the eudesmatic joint that ancestrally separated segments 7 and 8 corresponds to the paracoxal suture in Matsuda 1970^36^, and Kobayashi 2013^34^, 2017^35^, and 2018^60^. In I, L, morphological boundary between leg segments 7 and 8 of *Drosophila* kindly provided by Professor Yukimasa Kobayashi’s analyses of attachment points of muscles Iltpm4, 5, and 6, and Ipcm6 in the *Drosophila* thorax from Liu 2023, as well as the fact that no *Sp1* deletion phenotypes for body wall, pleura, or notum have been reported for *Drosophila*^93,177,187^. A, D from Pechmann 2011^79^, note that strong Dll RNAi also deletes walking leg 1. B, E, H, K from Bruce and Patel 2020^24^. C, F from Beerman 2001^88^. G, J from Konigsman 2017^82^. I, L from Estella 2010^93^. L shows an animal with a clone of cells generated during embryogenesis with a genomic deletion of both Sp1 (i.e. Sp6-9) and btd (Sp5, the less active paralog^187^), and Estella 2010 conclude that the remaining leg “consisted of only a small patch of residual leg tissue. When leg tissue was observed, it was invariably associated with non-mutant tissue, suggesting that these clones were generated slightly later than those samples in which no leg tissue remained.”. Abbreviations from source material: as, anterior spinneret; ch, chelicera; e, endite; L1-4, walking leg 1-4; pp, pedipalp; ps, posterior spinneret. See Fig. S2 for images without color.

In all arthropods examined, Iroquois genes are expressed strongly around proximal leg segment 8, with accessory expression domains, like the lateral dot and the tip of the claw, being variable in different arthropod lineages. However, the conserved expression configuration is *pnr* in a dorsal-medial stripe, which is apposed to a stripe of *iro*, with a second stripe of *iro* between leg segments 7 and 8.

## Discussion

The expression of *pnr*, *ara*, and *drm* reveals how the proximal leg segments of insects, crustaceans, and chelicerates correspond to each other, which supports the hypothesis that arachnids with 7 moveable leg segments possess an embryonic proximal 8^th^ leg segment that forms the lateral body wall in the adult. This is consistent with all available molecular, embryological, and morphological data in chelicerates, as follows.

To align the distal leg segments in insects, crustaceans, and chelicerates, we turn to the published molecular loss-of-function literature. Four leg patterning genes have been functionally tested in representatives for insects, crustaceans, and chelicerates: *Distal-less* (*Dll*), *Sp6-9*, *dachshund* (*dac),* and *homothorax* (*hth*). We note that the expression of these leg patterning genes is quite variable, both between arthropod phyla as well as over the embryonic development of each individual species^24,73^. For example, *Dll* is expressed in regions where it does not have a leg phenotype (early proximal leg^74,75^ and wing sensory hairs^76^), and conversely, *dac* is not expressed in regions where it does have a phenotype (leg segment 3, but presumably *dac* is expressed here earlier). Thus, the expression of these four genes is not as reliable as gene function when comparing such phylogenetically distant groups, and can confound interpretations. This is in contrast to the expression of *pnr*, *ara*, and *drm*, which – at the stage when leg joints are forming – are surprisingly well-conserved in their expression domains despite half a billion years of divergence.

We therefore focus on the loss-of-function data for *Dll*, *Sp6-9*, *dac*, and *hth* more than expression. Furthermore, given that RNAi produces a range of moderate to strong knockdowns, we focus on what appear to be strong phenotypes. For the following descriptions of leg segment deletion phenotypes, all interpretations are taken directly from the source material. For example, Setton and Sharma^77^ state that *Sp6-9* RNAi in spider results in “truncation of all remaining appendages up to the proximal-most leg segment, the coxa”, which in the present nomenclature is leg segment 7. In several cases, the respective authors were also contacted to confirm the interpretations here (Prashant Sharma 2025; Natascha Turetzek 2022; Mathias Pechmann, July 2022; Graeme Mardon, 2020; Carlos Estella 2018). Furthermore, the respective authors’ conclusions were visually confirmed from their original figures based on the shape of each leg segment. For example, spider leg segment 7 (coxa) is short and has a medial bulge called the coxal endite. This endite is especially pronounced on the pedipalp, which has the same number of leg segments as walking legs^78^ making the pedipalp endite a useful morphological marker for interpreting phenotypes. The interpretations from the source material are applied to the best of our ability in demarking leg segment boundaries in the figures reproduced here.

Based on the leg segment deletion phenotypes of *Dll*, *Sp6-9*, *hth*, and *dac,* the six distal leg segments of *Parhyale*, insects, and chelicerates (leg segments 1 – 6, counting from the distal claw) correspond to each other in a one-to-one fashion (Figs. 3, 4), as follows.

*Dll* RNAi in chelicerates deletes distal leg segments 1 – 5^79–81^ (Fig. 3D, and personal communication, Mathias Pechmann, July 2022), the same segments in which *Dll* is expressed when the embryonic leg joints become visible. The *Dll* RNAi phenotype is confirmed morphologically by the presence of the coxal endite in the leg stump, and molecularly by the expression in the stump terminus of *dac1*, a marker of leg segment 6^71,77,82–84^, the next most proximal leg segment. Thus, *Dll* is required for the development of leg segments 1 – 5 in chelicerates, a crustacean, and insects (Fig. 3A – F)^24,79,85–90^.

*Sp6-9* RNAi in chelicerates deletes distal leg segments 1 – 6^77,82^ (Fig. 3J). This is confirmed morphologically by the presence of the coxal endite on the pedipalp stump, indicating that leg segment 7 is still present, and molecularly by the lack of *dac1* expression, a marker of leg segment 6, in the stump, despite *dac1* expression in the ventral nerve cord^77,82^. Thus, *Sp6-9* is required for the development of leg segments 1 – 6 in chelicerates, a crustacean, and insects (Fig. 3G – L)^24,77,82,91–93^.

For *dac*, the available data is sparse and less clearcut, but suggests that *dac* patterns middle leg segments 3 – 5 (Fig. 4D-F). The strongest *dac* functional data for a chelicerate is in the harvestman *Phalangium opilio*^78^. Similar, but more moderate, effects are also seen in the spider, *Parasteatoda*^83^, where only one of the two *dac* paralogs gives an observable phenotype (Natascha Turetzek Zhang, personal communication). In the harvestman *Phalangium*, RNAi against the single copy of *dac* affects leg segments 3 – 5, such that segments 4 and 5 are deleted, and segment 3 is truncated (Fig. 4D). The crustacean, *Parhyale,* has two *dac* paralogs: *dac1* is more strongly expressed in a neural pattern and *dac2* is more strongly expressed in the medial leg^94^. Consistent with this expression difference, only *dac2* in *Parhyale* gives a discernible phenotype; *dac1* may have a neural phenotype but this was not observable from cuticle preps.

**Fig. 4.**
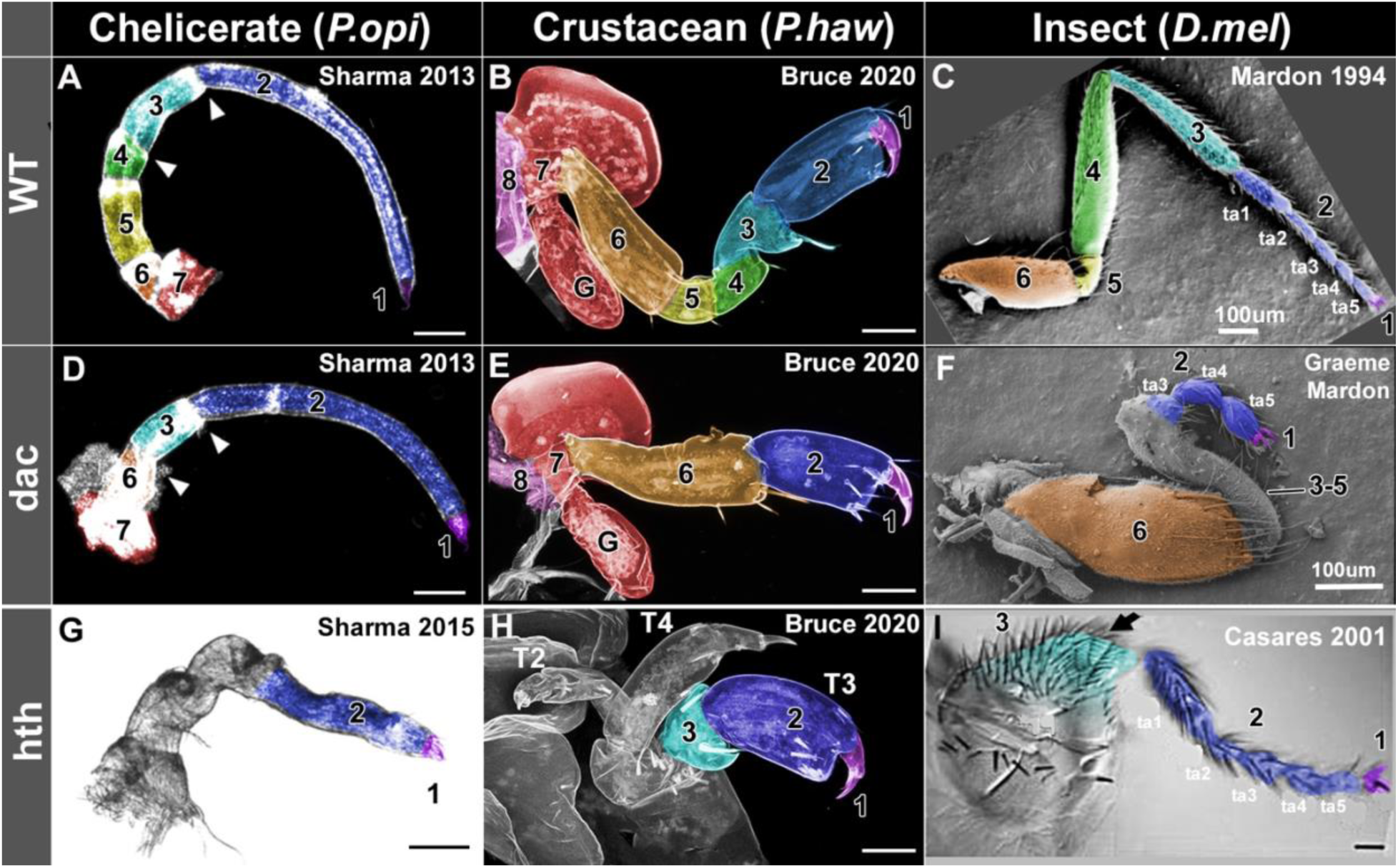
*dac* and *hth* function in arthropods. A-C, WT legs. D-F, *dac* loss of function. In chelicerate *Phalangium*, crustacean *Parhyale*, and insect *Drosophila*, a strong *dac2* phenotype affects leg segments 3 – 5. Note that the *dac* phenotype in *Drosophila* extends partway into leg segment 2, such that the proximal part (tarsomeres 1 – 2, ta1, ta2) are fused with medial leg segments 3 – 5 into a contiguous mass. However, tarsal bristles are present on the distal end of the fused mass (blue shading) and the joint with tarsomere 4 is intact, suggesting that tarsomere 3 is partially present. The coxa (leg segment 6) and tarsomeres 4 and 5 (distal end of leg segment 2) remain intact. G-I, *hth* loss of function. In *Parhyale*, genomic loss of *hth* deletes the proximal leg segments, leaving only the distal 2 leg segments intact. In *Phalangium*, reduction of *hth* mRNA shortens and fuses the proximal leg segments, leaving the distal segments unaffected. Arrow in I, transverse row bristles, demarcating the distal end of leg segment 3. Scale bars 50µm except where indicated. A, D, G from Sharma 2013^78^, 2015^97^. B, E, H from Bruce and Patel 2020^24^. C from Mardon 1994^99^. F, dac^4^/dac^4^ from Graeme Mardon, unpublished. Note this panel is zoomed in to 2x the magnification compared to the WT in C so that the reader may confirm the details of the phenotype. I from Casares 2001^98^. Abbreviations: T2-4, third thoracic legs. See Fig. S3 for this figure without color.

This is similar to the situation in spider, where only one of the two *dac* paralogs gives an observable phenotype, which suggests that one *dac* paralog in *Parasteatoda* and *Parhyale* may retain the medial leg patterning while the other controls other aspects of patterning. In *Parhyale*, CRISPR-Cas9 knockout of *dac2* completely deletes leg segments 3 – 5, such that leg segment 2 connects directly to 6^24^, even though *dac* is not clearly expressed in segment 3 at the stages examined^24^. In the insect *Drosophila*, loss of *dac* also affects leg segments 3 – 5 (as well as the proximal part of 2), which are shorter and fused into a contiguous mass of tissue^95^. RNAi against *dac* in the insect *Oncopeltus* deletes leg segment 3 and shortens leg segment 4^89^, and RNAi against *dac* in *Tribolium* deletes leg segments 3 and 4, as well as the proximal part of 2^96^. Thus, *dac* appears to be required for the development of leg segments 3 – 5 in a chelicerate, a crustacean, and insects (Fig. 4A – F), but insect *dac* function appears to have shifted slightly distally^24,78,83,89,95,96,99^. Strong CRISPR-Cas9 phenotypes in chelicerates, myriapods, other crustaceans, and non-*Drosophila* insects would clarify this ambiguity.

For *hth*, the only chelicerate for which there is functional data is the harvestman, *Phalangium opilio*. In *Phalangium*, RNAi against *hth* shortens and fuses the proximal leg segments, leaving the distal segments unaffected^97^. Given that the most severely affected embryos died before hatching, their wrinkled cuticle obscured the joints, making it difficult to discern with high certainty how many distal leg segments are unaffected. However, in the normal-looking distal part of the leg, leg segment 1 and the joint its shares with leg segment 2 are present, so the distal half of leg segment 2 appears unaffected. Thus, *hth* is required for the development of the proximal leg segments and leaves distal leg segments 1 and part of intact 2 in arachnids as well as *Parhyale* and insects (Fig. 4H – J)^24,89,95,97–100^.

The functional data for *Dll*, *Sp6-9*, *dac*, and *hth*, together with the conserved expression patterns of *pnr*, *ara*, and *drm*, support a model wherein living arthropods ancestrally had 8 leg segments, and that the reduced number of canonical leg segments in chelicerates, myriapods, non-insect crustaceans, and insects appears to be due to the conversion of proximal embryonic leg segments into lateral body wall in the adult.

The molecular data discussed here agrees with over a century of morphological, embryological, and paleontological observations suggesting that arthropods ancestrally had 8 leg segments. First, living arthropods have no more than 8 true (eudesmatic) leg segments posterior to the first antenna^2^; early-branching chelicerates^48,49,104^ often have 8 leg segments^4^; many crustaceans have an additional proximal leg segment, known as the precoxa^2,3,10^, corresponding to leg segment 8 here; and the fossil ancestors of all living arthropods had 8 leg segments^46^.

Furthermore, in arthropods with fewer than 8 moveable leg segments, the embryonic leg base is observed to broaden and expand to form the adult lateral body wall^34,35,37,47,105,106^, suggesting that the arthropod ancestor had more leg segments than the present number of moveable leg segments would indicate. Similarly, pleurites (hard plates floating in the lateral body wall to which muscles attach, often surrounded by soft cuticle) are found only in arthropods with fewer than 8 free leg segments (Fig. 6b), including insects^8,34,35,51,107^, decapods (crustaceans)^1,25,108,109^, millipedes^28,110^, centipedes^1^, and pauropods^105^ (myriapods), and horseshoe crabs (chelicerate)^1^.

Notably, pleurites appear to be derived from the embryonic leg base^1,34,105,107^, thus accounting for the missing ancestral leg segments in these lineages. These morphological, embryological, and fossil observations have led previous authors to a similar conclusion within each arthropod lineage^4,8,10,46,51^. These are here combined with the added insight of modern molecular phylogenies^48,49,111,112^ and gene expression and function data to infer that all arthropods ancestrally had 8 leg segments that correspond to each other in a one-to-one fashion (Fig. 6). For example, the insect coxa, crustacean basis, and spider trochanter correspond to each other, such that a simpler term for these would be “leg segment 6”, while the insect trochanter, crustacean ischium, and spider femur could be referred to as leg segment 5.

The leg model presented here also explains another peculiarity: a vestigial leg joint within the adult insect lateral body wall. Arthropod leg joints can be defined molecularly by the expression of joint patterning genes like *odd-skipped* family genes^53,68,69,101,102^ and *Serrate*^37,103^; and morphologically by the presence of a thin, flexible line of cuticle (called a suture) with muscle insertions^2,36^. The vestigial leg joint present in the lateral body wall of adult insects satisfies both of these criteria: *odd-skipped* genes and *Serrate* are expressed in a ring here in embryos, and this region forms a morphological suture known as the paracoxal suture in the adults of many insect lineages onto which muscles insert^35–37^, which represents the vestigial joint between leg segments 7 and 8 that forms the lateral body wall of the adult insect (Fig. 5b).

**Fig. 5.**
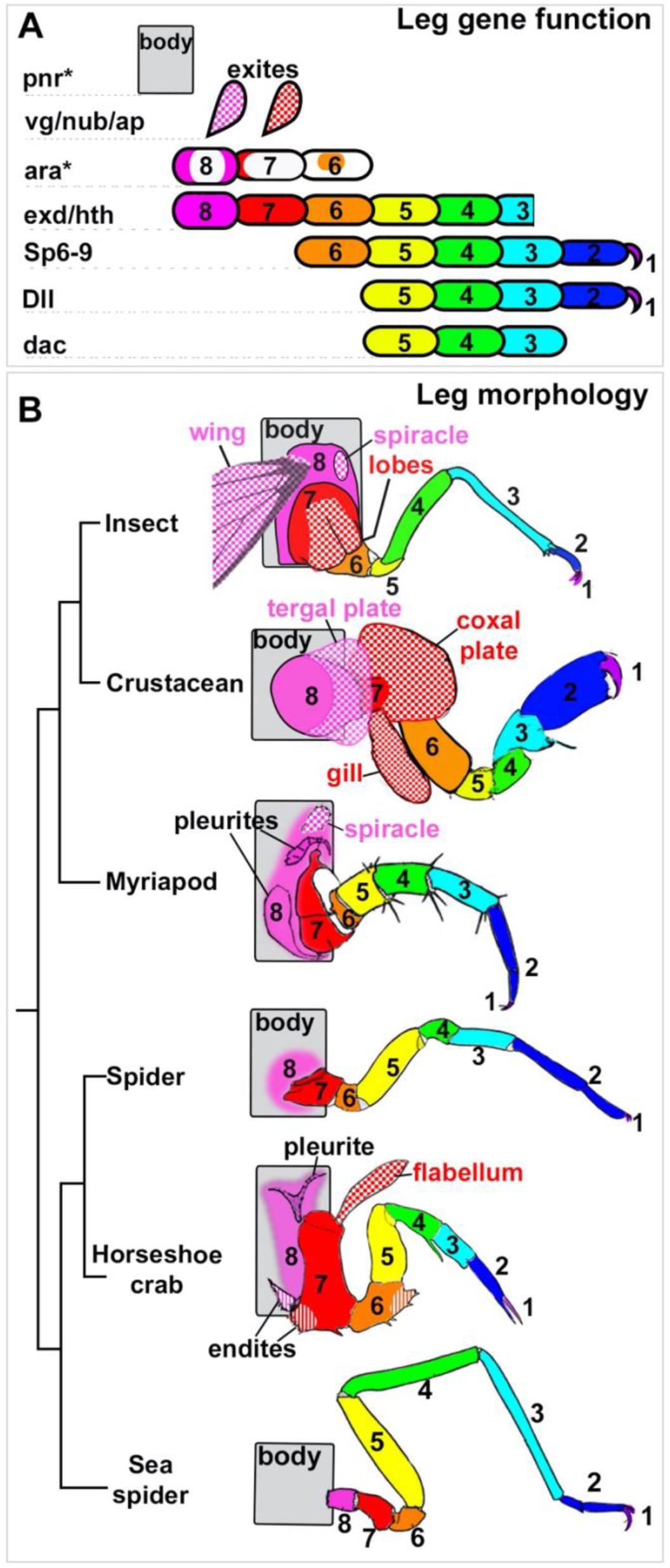
Proposed model for how all arthropod leg segments correspond to one another. A. Schematic of the leg segments and structures patterned by each gene. B. Morphology and homologies of arthropod leg segments based on leg gene function in insects, *Parhyale*, and chelicerates. Colors and patterns indicate proposed homologies. Exites (checker pattern); endites (stripe pattern). Drawings in B modified from Snodgrass 1952.

The differences in the canonical leg segment numbers in the four groups of arthropods are proposed to be due to the variable conversion of proximal leg segments into the body wall, likely to support more of the body weight on the legs during the transition from swimming to walking and from aquatic to terrestrial lifestyles. Chelicerates with 7 free leg segments, like spiders, have converted one leg segment into the body wall (leg segment 8), whereas chelicerates with 8 free leg segments, like sea spiders, camel spiders, and red velvet mites, appear to retain the proposed ancestral number of 8 free and mobile leg segments^4^. Myriapods with 6 or 7 free leg segments^1,2,7,28,52,110,113,114^ appear to have converted 2 or 1 leg segments into the body wall, respectively^105^. Insects, with 6 free leg segments, have converted two proximal leg segments into the body wall (leg segments 7 and 8). Crustaceans have variably converted the leg base into the lateral body wall. Many early-branching crustaceans^111,112^, like branchiurids, mystacocarids and myodocope ostracods, retain a free and mobile leg segment 8 (precoxa)^2,3,10,115–118^. In decapods like prawn and crab, the embryonic/ancestral precoxa forms the adult lateral body wall, such that the pleurobranch gills (and probably arthrobranch gills) that are carried on the adult lateral body wall^23,119^ are derived from the embryonic precoxa^10,108,109,119,120^. In amphipods like *Parhyale*, the embryonic precoxa forms much of the lateral body wall, but is obscured by the tergal and coxal plates^24^. However, the precoxa is revealed when wing genes like *vestigial* are knocked out, which deletes the tergal and coxal plates, leaving behind just the exposed, protruding precoxa^16^. Terrestrial isopods with 6 free leg segments have, like insects, converted two proximal leg segments into the body wall (leg segments 7 and 8), an evolutionary sequence which is easier to appreciate in this lineage because aquatic isopods still retain a free and mobile leg segment 7^121–124^. Mantis shrimp possess a free and mobile precoxa but only on the posterior three thoracic legs; the powerful raptorial legs do not have a free precoxa^3,125–127^, which may have converted the precoxa into the body wall in order to provide a more robust base from which to exert force. Cladoceran crustaceans like the water flea *Daphnia* have partially converted the precoxa into the lateral body wall^10^.

Insect embryologists since the late 1800s, working in several major clades of insects, have observed that the embryonic leg base flattens and expands to form the lateral body wall of the adult^8,34,36,37,47,54,128–131^. This has become known as the insect subcoxa theory, likely familiar to any entomology student. The analysis presented here suggests that this same morphogenetic sequence happens throughout arthropods, meaning that the insect subcoxa theory should more generally be called the arthropod subcoxa theory.

### A system to decipher the origins and relationships of any arthropod outgrowth

In the model presented here, the function of the five leg patterning genes in conjunction with the expression of the proximal patterning genes *pnr* and *ara* and the joint marker *drm* can be used to infer the homologies of structures along the proximal-distal axis in the same way that the well-known “Hox code” has been used to infer homologies along the anterior-posterior axis. Notably, given the highly conserved expression patterns of *pnr*, *ara*, and *drm* across arthropods, and the ease and low cost of in situ Hybridization Chain Reaction (HCR)^132,133^ expression experiments, the homologies of any and every long-debated arthropod ectodermal structure can now be elucidated with a single in situ HCR experiment using just three or four genes. For example, a simple four-gene HCR of *pnr, ara, drm*, and a wing gene like *vestigial* could answer definitively whether beetle horns and treehopper helmets are wing serial homologs that are carried on embryonic/ancestral leg segment 8. Similarly, a simple three-gene HCR of *pnr*, *ara*, and *drm* could answer definitely whether all “entomostraca”-like crustaceans truly lack abdominal legs^134^, or have simply truncated them as insects have^135^. Additionally, if insects have 6 free and 2 converted leg segments, such that *odd-skipped* genes are expressed in a stripe at the vestigial joint between segments 7 and 8 that form the lateral body wall, then other arthropods with 6 free and 2 converted leg segments should also express *odd-skipped* genes in a stripe at this vestigial joint, such as isopod crustaceans, and symphylan and pauropod myriapods^2,52,105,113,114,121,122^. Furthermore, the expression of *pnr* and *ara* has thus far only been examined in arthropods that have partially or completely converted leg segment 8 into the lateral body wall. The expression of these genes should therefore be examined in arthropods where leg segment 8 is still free and mobile, such as sea spiders (Fig. 5), camel spiders (solifugids), red velvet (trombidiid) mites, and mantis shrimp (leg segment 8/precoxa is free^125–127^, but the distal leg segment number is reduced; however, leg segment 6/basipod is easily identifiable morphologically by the presence of the exopod^3,125,126^). If this model is accurate, then *pnr* should be expressed in the true body wall (perhaps only dorsally but not ventrally), *ara* should bracket the moveable 8^th^ leg segment proximally and distally, and *odd-skipped* genes should only be expressed where two leg segments meet, but not where the proximal 8^th^ leg segment meets the body wall. Similarly, a prediction of this model is that lateral outgrowths like wings, plates, gills, carapaces, and horns should only form on leg-derived tissue (either free and moveable leg segments, or on the leg-derived body wall), but not on the non-leg derived “true” body wall. A related prediction then is that arthropods that retain 8 moveable leg segments, like sea spiders, camel spiders (solifugids), red velvet (trombidiid) mites, and mantis shrimp, should not have outgrowths of their body wall, which seems to be the case. While the mantis shrimp may seem to contradict this prediction in that it has both a free precoxa and a carapace, it in fact proves the point: the legs that possess a moveable precoxa (the three posterior thoracic legs) do not form the carapace; the carapace is formed by the anterior legs that do not have a free precoxa^3,125,136^.

### Determining the leg segment affinities of chelicerae

An example of how this leg coordinate system can be put to work with a simple HCR experiment is the chelicerae (anterior-most legs of chelicerates). Chelicerae are the namesake of the chelicerates, and represent the appendages of the deuterocerebrum that correspond to the antennae of insects, the first antennae (antennules) of crustaceans^137–140^, and perhaps the great appendages of fossil megacheiran arthropods^141^. In spiders, the chelicerae form fangs, but in sea spiders, horseshoe crabs, scorpions, and mites, they form pincer claws. Chelicerae have been proposed to have been 3-segmented in the chelicerate ancestor, but that the proximal segment was lost in more derived spiders with 2-segmented chelicera^142,143^. The expression of *pnr, ara, drm,* and *Dll* in *Acanthoscurria*, an early-branching spider, supports this hypothesis in that the proximal-most leg segment 8 appears to form the body wall of the head, as follows (Fig. S3). *pnr* is expressed dorsally around the head lobes, likely marking the non-leg-derived ancestral body wall. *ara* is expressed in the same three domains as the other legs of *Acanthoscurria*: the dorsal armband that follows the *pnr* domain, the second armband, and the distal claw domain in the fang. *drm* is expressed in two bands, indicating the boundaries of three leg segments. But which of the eight possible leg segments in particular? The proximal-most segment likely corresponds to leg segment 8, because it is bracketed by the dorsal *ara* armband that follows *pnr* and the ventral *ara* armband, which is also adjacent to the *drm* band, indicating the distal end of the segment. The distal-most segment of the chelicera likely corresponds to leg segment 1 (the claw) because of the distal, sock-like expression of *Dll* as well as *ara* in the fang. The identity of the medial segment is more ambiguous. It expresses *Dll*, and therefore could correspond to leg segments 2, 3, 4, or 5. However, a comparison of *dac* expression in spiders narrows the possibilities because *dac* is co-expressed with *Dll* only in leg segment 5 (femur)^73,83,84,143^.

### Myriapod legs

Unfortunately, no gene functional data is available for myriapods, and in myriapod embryos, the non-specific background staining becomes too high after the early leg bud stages when the leg joints form, which precludes relating leg gene expression domains to specific morphological leg segments. However, the available gene expression data for early leg patterning is consistent with the model presented here.

In all arthropods including myriapods, *Dll* is expressed in distal sock-like domains of the mouthparts and legs^144–146^, corresponding to leg segments 1-5. Consistent with the model presented here, the mandible of mandibulates (myriapods, crustaceans, and insects) appears to lack segments 1-5 (the telopod), and consist of only the two proximal-most leg segments, 7 and 8^68,77,145–147^, with segment 8 latter forming the head lobes and conglomerated walls of the head, and segment 7 forming the moveable part of the mandible. For example, *Dll* is expressed only in the tooth-like structure of the mandible, and appears to have a sensory role rather than patterning the actual leg-segments of the mandible^24,68,94,145–147^. Similarly, in all arthropods including myriapods, *exd* and *hth* are expressed in proximal domains, including in the mandible^144,148^, whereas *dac* is expressed in a medial domain in non-truncated legs, but in the distal tip of the mandible and maxilla^144^. *dac* expression in the mandible likely represents endite patterning^106,144^, similar to *Dll* expression here, while dac expression in the maxilla may correspond to endite patterning or to leg segment 5 in these distally truncated leg types^94,149^.

Furthermore, the expression domains of axial patterning genes, which also pattern legs, are highly conserved across all arthropods, including myriapods^150,151^. For example, *engrailed* and *hedgehog* are expressed in the posterior of each leg, while *cubitus interruptus* is expressed in the anterior^152–154^. Similarly, *wingless* is expressed in a ventral stripe in each leg, while *decapentaplegic* and *optomotor blind* are expressed dorsally in each leg^153,155,156^.

No data is available for the expression of *pnr* or any of the Iroquois genes like *ara* in any myriapods, which would facilitate the molecular identification of the proximal leg segments.

However, the morphological and embryological data suggests that in myriapods, like insects, many crustaceans, and spiders, the embryonic leg base forms the adult body wall^2,105,110^. In embryos of a myriapod representative, the symphylan *Hanseniella*, the proximal part of the developing leg (“limb base”) is observed to broaden and flatten to form the adult lateral body wall^105^ . Incorporation of proximal leg segments into the myriapod body wall would bring their leg segment count to 8, in agreement with other arthropods.

A model and simple 3-gene test to decipher the origins and relationship of any ectodermal structure in any arthropod opens up a powerful system for studying the origins of novel structures, the evolvability of morphology and gene networks across vast phylogenetic distances, and the convergent evolution of shared ancestral developmental fields.

## Materials and Methods

### Embryo fixation

*Acanthoscurria* embryos were fixed overnight in 5% formaldehyde/PBS and heptane. *Tribolium* embryos were fixed 30 min in 4% formaldehyde/PEM and heptane. *Parhyale* embryos fixed 40 min in 4% PFA in or artificial seawater. All embryos then dehydrated into 100% methanol. All embryos stored at -20°C.

In situ HCR version 3.0

In situs were performed as in Bruce et. al. 2021^133^. cDNA sequences were submitted to Molecular Instruments^132^, and the probe sets are available from the company, as below. cDNA sequences for Acanthoscurria *pnr, ara, Dll*, and *drm* Supplemental information. Probe sequences for the three *pannier* paralogs ordered as o-pools from IDT in Supplemental information.

| Species | Gene | Accession number | Probe set numbers from Molecular Instruments |
| --- | --- | --- | --- |
| *Acanthoscurria geniculata* | *araucan* | - | PRE883 |
| *Acanthoscurria geniculata* | *Distalless* | - | PRE884 |
| *Acanthoscurria geniculata* | *drumstick* | - | PRG135 |
| *Tribolium castaneum* | *pannier* | XM_008202266.2 | PRD736 |
| *Tribolium castaneum* | *araucan* | XM_008194186.2 | PRD737 |
| *Tribolium castaneum* | *Distalless* | NM_001039439.2 | PRD738 |
| *Parhyale hawaiiensis* | *pannier* | MT103930.1 | PRD223 |
| *Parhyale hawaiiensis* | *araucan* | MT103931.1 | PRD221 |
| *Parhyale hawaiiensis* | *Distalless* | FJ617569.1 | PRD002 |

### Imaging

Embryos imaged with Zeiss LSM880 confocal. Image processing done with Fiji-ImageJ. Fiji “Image Calculator > Subtract” method was used to remove high background from yolk autofluorescence. Figures processed using Adobe Photoshop 2020.

### Gene orthology

OrthoFinder and gene phylogenies were used to determine orthology of Acanthoscurria *pannier*, *araucan*, and *drumstick*. *Acanthoscurria* transcriptome, kindly provided by Mathias Pechman, and *Parhyale* transcriptome from Sun 2022^157^ were each first converted to proteomes using Transcdecoder.LongestORFs^158^, then Transdecoder.Predict with the single_best_orf option and blastp options using the Drosophila Reference Sequences proteome (NCBI), Tribolium Reference Sequences proteome (NCBI) to identify the single best ORF for each transcript. The resulting *Acanthoscurria* and *Parhyale* proteomes were then used with OrthoFinder against the above proteomes for *Drosophila* and *Tribolium* (which each contain the specific sequences of *pnr*, *ara*, and *drm* that have been previously verified by in situ to give the expected expression pattern), to identify *pnr*, *ara*, and *drm* in *Acanthoscurria* and *Parhyale*.

The resulting Acanthoscurria proteome was also used to generate phylogenies for pnr and ara. For ara, the following sequences with accession numbers were used to generate sequence alignments, with the closely related Mohawk gene^159^ as the outgroup:

NP_808263.2_Mohawk_Mus, XP_001811513.2_Mohawk_Tribolium, NP_608502.1_CG11617_Mohawk_Drosophila, XP_008192408.1_araucan_Tribolium, XP_008192405.1 araucan_Tribolium, NP_001163432.1_araucan_Drosophila, KYB28757.1_araucan_Tribolium, KYB28756.1_araucan_Tribolium, XP_008192400.1 araucan_Tribolium, AAT11862.1_iroquois_Tribolium, AAB53640.1_mirror_Drosophila, XP_971676.1 caupolican_Tribolium, KZS08873.1_caup_Daphnia_magna, NP_524046.2_caup_Drosophila, XP_971676.1 caup_Tribolium, XP_016770700.1 caup_Apis, KFM58187.1_caup_Stegodyphus, NP_524045.2_araucan_Drosophila, NP_001163432.1_araucan_Drosophila, XP_015182907.1 araucan_Polistes_dominula, XP_018911022.1 ara_Bemisia_tabaci, JAG00978.1_araucan_Lygus_hesperus, ODM99984.1_ara_Orchesella_cincta, XP_008192408.1 araucan_Tribolium, XP_008192405.1_araucan_Tribolium, XP_008192400.1_araucan_Tribolium, EKC34186.1_araucan_Crassostrea, NP_524047.2_mirror_Drosophila, EHJ65533.1_mirror_Danaus, JAB79499.1_mirror_Ixodes, JAB72752.1_mirror_Ixodes, XP_015907356.1_caup_Parasteatoda, EEC00349.1_IRX_Ixodes, XP_015782374.1_IRX-3_Tetranychus, KFM58187.1_caupolican_Stegodyphus, XP_002400564.1_IRX_Ixodes_scapularis, KZS08873.1_caup_Daphnia_magna, XP_021002308.1_IRX-6_Parasteatoda, XP_021002310.1_IRX-6_Parasteatoda, XP_021002309.1_IRX-6_Parasteatoda, KZS08871.1_IRX-1_Daphnia_magna, Parhyale.ara.phaw_50.282976aG352.1

For pnr, the following sequences with accession numbers were used to generate sequence alignments, with *Caenorhabditis elegans* elt-1 as the outgroup:

Hyalella_grn_XP_018013183.1, Parhyale_grain_phaw_50.000203bG892.1, Trinorchestia_grn_KAF2359614.1, Acanthoscurria_DN76703_c4_g1_i1, Acanthoscurria_DN74902_c0_g1_i1, Acanthoscurria_DN81267_c7_g2_i1, Stegodyphus_GATA4_KFM82098.1, Parasteatoda_GATA4_XP_015929395.1, Acanthoscurria_DN76525_c1_g1_i1, Euperipatoides_GATA123_CUW78639.1, Limulus_grn_XP_022242144.1, Limulus_GATA2_XP_022240403.1, Limulus_GATA2_XP_013773452.2, Euperipatoides_GATA456_CUW78640.1, Stegodyphus_GATA-4_KFM82100.1, Stegodyphus_pnr_XP_035228380.1, C_elegans_elt-1_NP_001368482.1, Tigriopus_grn_TRY70078.1, Stegodyphus_grn_XP_035208929.1, Parasteatoda_grn_XP_021003356.1, Trinorchestia_pnr_KAF2355501.1, D.magna_grn_XP_032783972.1, Eurytemora_grn_XP_023338271.1, Penaeus_grn_XP_027213737.1, Penaeus_grn_XP_027213731.1, Parhyale_grain_comp168149_c1_seq6, Tribolium_grain_NP_001158260.1, Drosophila_grain_NP_001262366.1, Centruroides_GATA-4_XP_023233756.1, Eurytemora_grn_XP_023340853.1, Penaeus_pnr_XP_027229190.1, Acanthoscurria_DN55779_c1_g1_i1, Hyalella_pnr_XP_018008986.1, Parhyale_pannier_comp167878_c4_seq4, Ixodes_GATA-5_XP_029839700.1, Limulus_pnr_XP_013775036.1, Limulus_GATA-4_XP_022242593.1, Ixodes_pnr_XP_029842940.1, Parasteatoda_GATA-A_BBD75256.1, Parasteatoda_LOC107456523_XP_015929888.1, Parasteatoda_LOC107456523_XP_015929890.1, Ixodes_GATA-5_XP_029840357.1, Acanthoscurria_DN77434_c10_g2_i1, Acanthoscurria_DN77434_c10_g2_i2, Centruroides_pnr_XP_023233757.1, Centruroides_grn_XP_023227752.1, Ixodes_grn_EEC05487.1, Parasteatoda_LOC107439679_XP_015907840.1, Parasteatoda_GATA-B_BBD75257.1, Acanthoscurria_DN80076_c8_g1_i4, Limulus_GATA-B_XP_022249933.1, Acanthoscurria_DN78099_c0_g1_i1, Acanthoscurria_DN77604_c0_g1_i1, Centruroides_GATA-4_XP_023228374.1, Stegodyphus_GATA-5_KFM72207.1, Stegodyphus_GATA-A_KFM57600.1, Parasteatoda_pnr_XP_015929399.1, D.pulex_pnr_EFX78972.1, D.magna_GATA_KZS08761.1, Bombyx_pnr_QGA69977.1, Tribolium_serpent_XP_008200495.1, Tigriopus_pnr_TRY71481.1, Glomeris_pannier, Nasonia_pnr_XP_001607848.1, Tribolium_pannier_XP_008200486.1, Ontho.sag_pnr_QDE10346.1, Drosophila_pannier_NP_001262620.1, Apis_pnr_XP_001121210.2, Ontho.taur_pnr_XP_022901096.1 Sequence alignments were generated using clustal^160^, mafft^161^, and muscle^162^. Best model was selected using ProTest^163^. Trees were then generated in Geneious^164^ with PHYML^165^ and MrBayes^166^ using the recommended model from ProtTest. Trees visualized with TreeView^167^

## Data and materials availability

All data is available in the main text or the supplementary materials.

## Acknowledgments

HSB thanks Nipam H. Patel for reagents, space, and equipment for this work. HSB thanks Carsten Wolff, Rachel Thayer, Joanna Wolfe, Gregory Edgecomb, and Frank W. Smith for helpful feedback. Fixed *Acanthoscurria* embryos in methanol, *Acanthoscurria* transcriptome, and unlabeled images of Dll RNAi spider embryos were kindly provided by Mathias Pechmann (Institute for Zoology/Developmental Biology, Biocenter, University of Cologne, Germany). HSB thanks Professor Yukimasa Kobayashi for analysis of *Drosophila* musculature to determine the location of the paracoxal suture boundary between leg segment 7 and 8 (Kobayashi’s subcoxa 2 and 1, respectively). HSB thanks Aaron Vose for computational assistance.

## Author contributions

HSB conceived, designed, and performed experiments, and wrote manuscript.

## Acanthoscurria sequences and probes

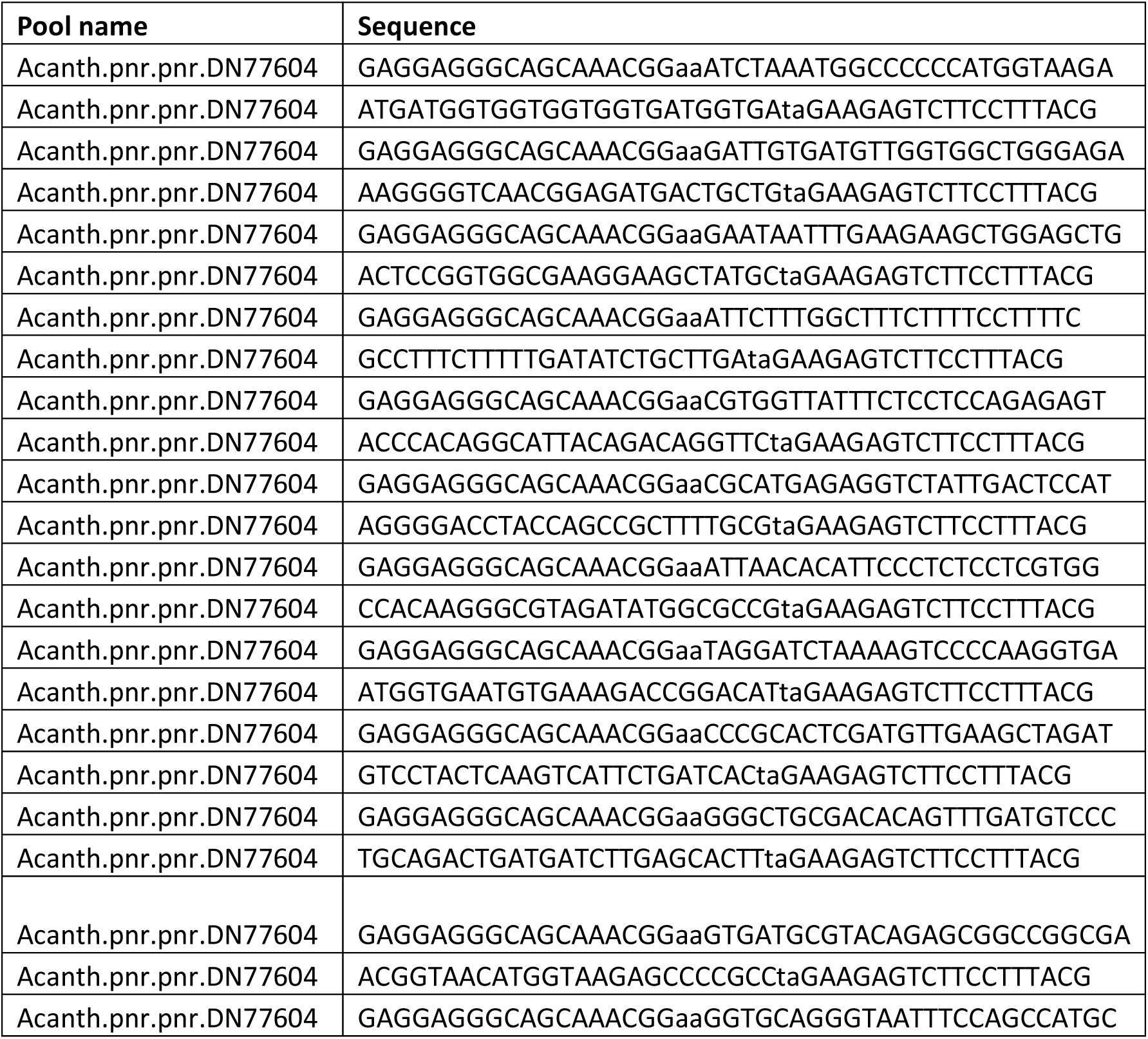
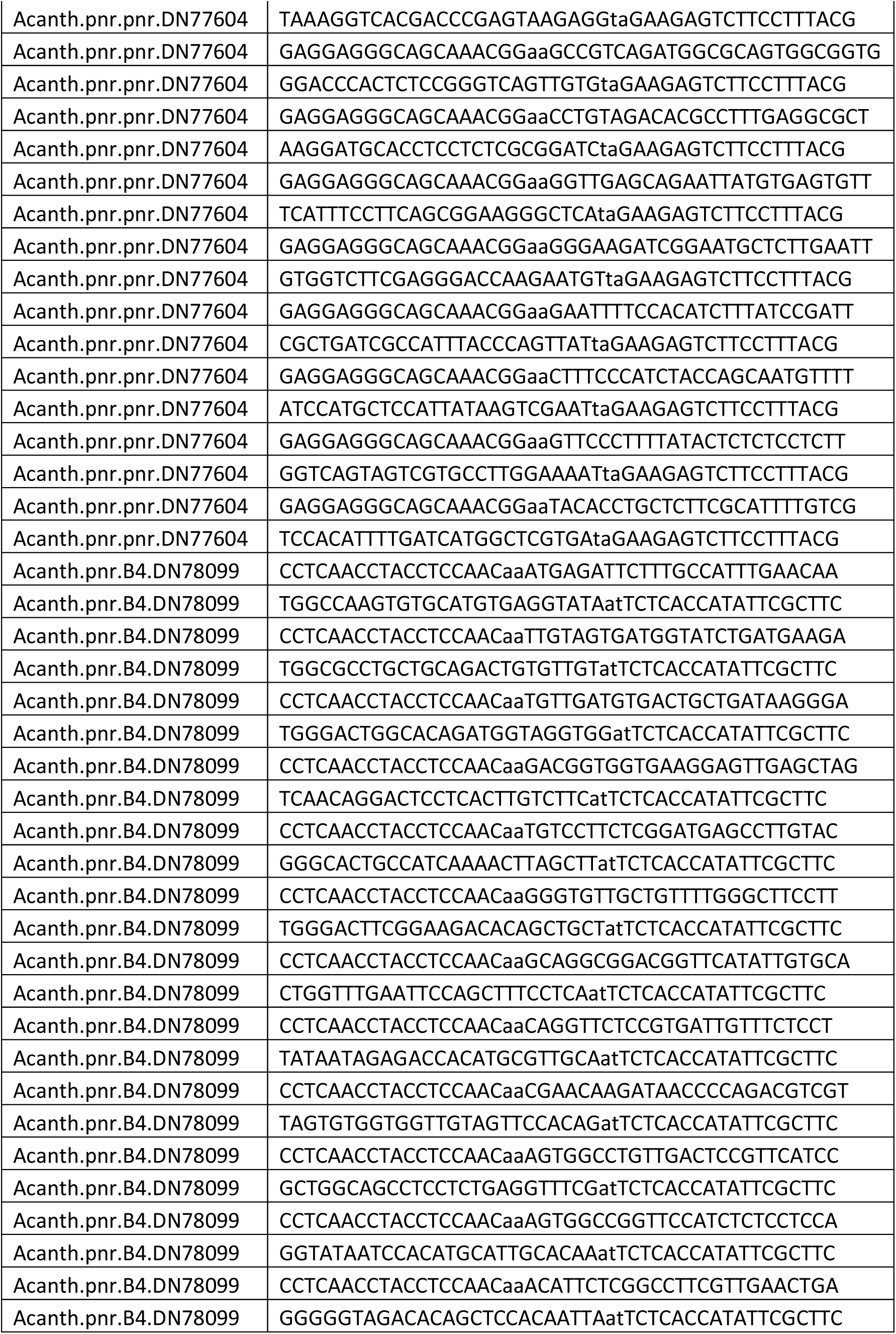
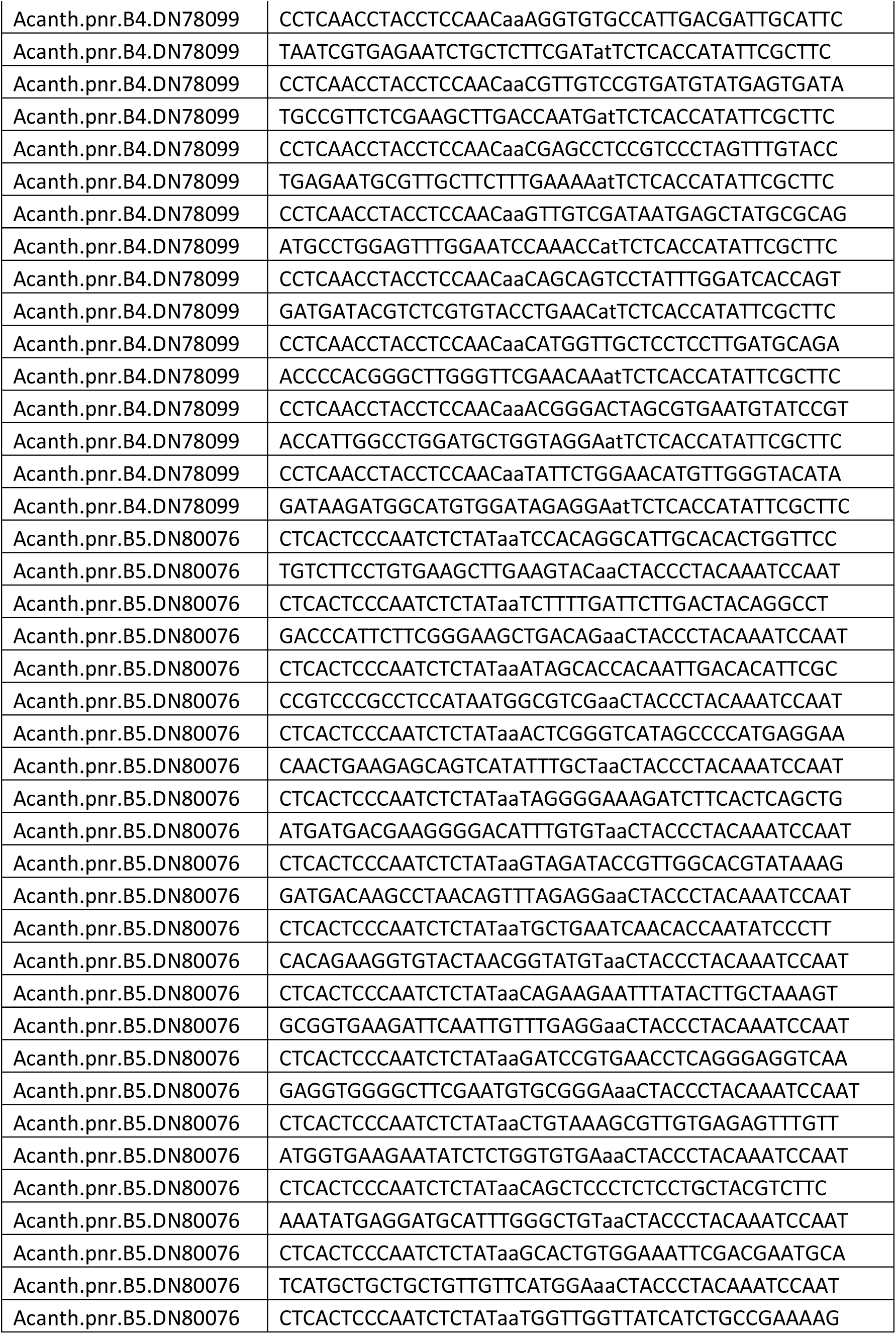
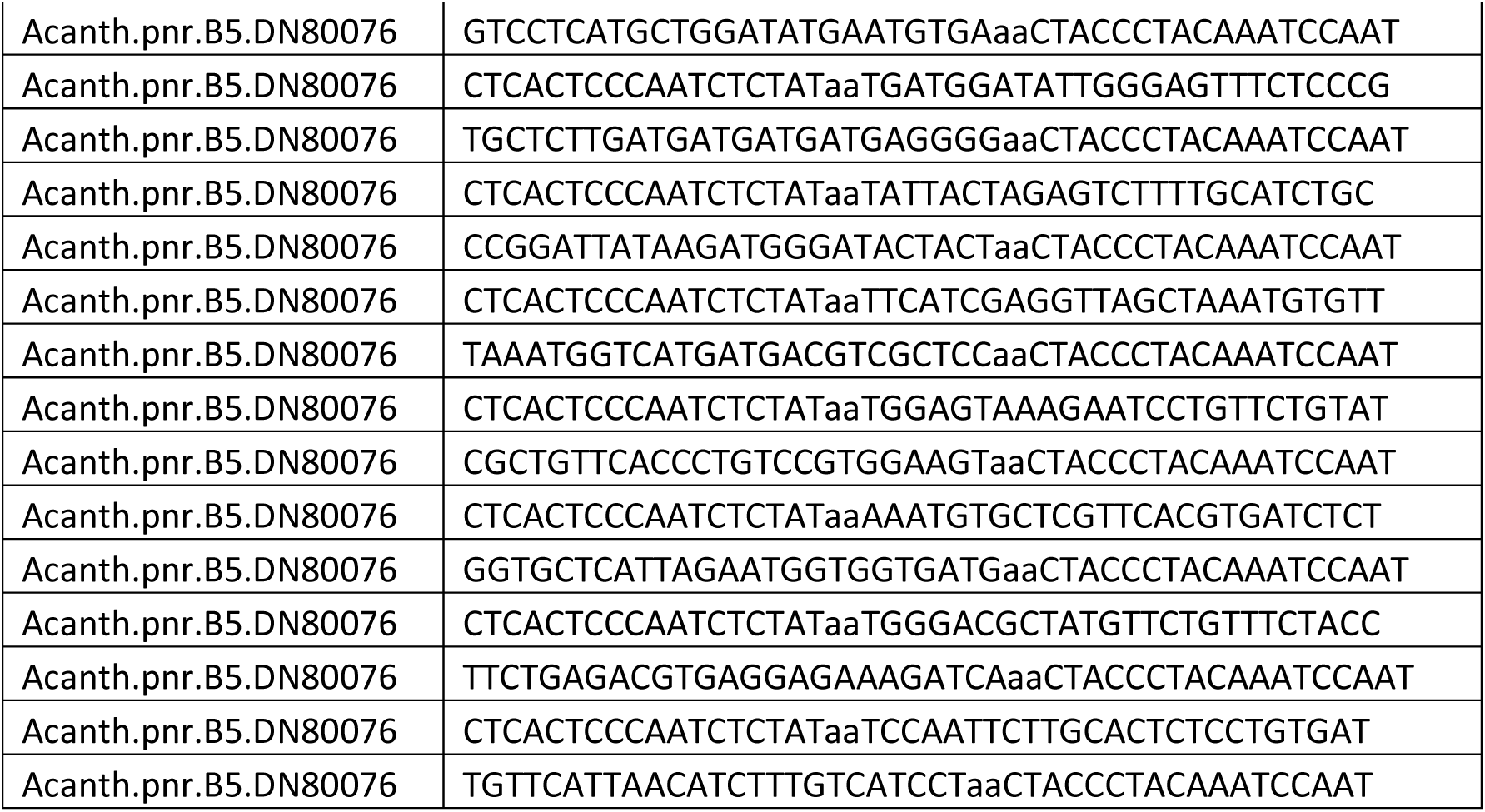

## Acanthoscurria sequences

### pannier

>pannier_TRINITY_DN77604_c0_g1_i1 CAAAATGATGAAAATTCTCACGTTTCGCTTCATGATATTCATTCTTGGAGCAATCACGAT AGTGCTGTAACAAGCCATTCTTGGGCAGATCATCCTTCTTCTCATAGTTCTAGTGTTCCA AAAACATCATCATGGCCTGCGAAGCAACATCCAACTTCGTTGCATCAAGGTGTTGATCTT CTGTGTCACGCCTGGGAAAAAACAAACGAGGAAAATCCAGAGCATTCACCTGGAGACTGG ATGAAAAATGAAGTGACGACAAAATGCGAAGAGCAGGTGTACTTCACGAGCCATGATCAA AATGTGGACGTAAGAGGAGAGAGTATAAAAGGGAACAAATTTTCCAAGGCACGACTACTG ACCAACAAAACATTGCTGGTAGATGGGAAAGTAATTCGACTTATAATGGAGCATGGATCA GAATCGGATAAAGATGTGGAAAATTCAGATAACTGGGTAAATGGCGATCAGCGTACAATT CAAGAGCATTCCGATCTTCCCTCACATTCTTGGTCCCTCGAAGACCACGCCCATATCAAC GGTATAGTGTCACATTCTGCGCCTGAATCGTCTAATAACCCCTGGGAACACTCACATAAT TCTGCTCAACCAATGAGCCCTTCCGCTGAAGGAAATGAAACACACCCTAGTGAGATAAAC ATCAACCAATCGGCCTCTGAAACCCCACCCACCAACAGCGCCTCAAAGGCGTGTCTACAG GGTGATCCGCGAGAGGAGGTGCATCCTTCCGAGCAGCGGAATCACTCCAGCGTAGTTCAC TTGCGATCGTCACAACAAGAACCCGACACCGCCACTGCGCCATCTGACGGCGGCACAACT GACCCGGAGAGTGGGTCCGACCACCCTAACAGTGTCCCAGTCATGGTCAGCACGTTTGGC AGCATGGCTCCATACAGCATGGCTGGAAATTACCCTGCACCATCCTCTTACTCGGGTCGT GACCTTTATTCTGGGAAACCATCTCCCGGCTACGGACATTCTGTGGACACCTCTTCTCCT ACAACTTCGCCGGCCGCTCTGTACGCATCACCGGGCGGGGCTCTTACCATGTTACCGTAT GTGACGGGTCAGTCAATGAGCCATCAATCCGTGGTCACCCAGGGCAGTCACGCATGGGGA CATCAAACTGTGTCGCAGCCCCCAAGTGCTCAAGATCATCAGTCTGCAAGTACATACGGA ATTTCTGCTTTGTCGTCAGGATTGAACTTGGCTCAAAGTTCTCTTAATCTAGCTTCAACA TCGAGTGCGGGATGTGATCAGAATGACTTGAGTAGGACAGGTGGATTTTCTACTTTCGGA TCCACTGGACATGGTTATCTTCGTCCGGACATGACGTCACCTTGGGGACTTTTAGATCCT ACAATGTCCGGTCTTTCACATTCACCATATTGTACAGATGGAATGGGACATCATTTAGGA CAAGATCGTGGGGACTATTTTGGCATCCACGAGGAGAGGGAATGTGTTAATTGCGGCGCC

ATATCTACGCCCTTGTGGCGGCGAGACGGGACCGGCCACTACCTATGCAACGCCTGTGGA CTCTACAGCAAGATGAATGGAGTCAATAGACCTCTCATGCGACCGCAAAAGCGGCTGGTA GGTCCCCTAGTGACAGCGAATCGAAGAGCCGGTCAGATCTGCACAAACTGTGGAACAACC ACGACTACTCTCTGGAGGAGAAATAACCACGGTGAACCTGTCTGTAATGCCTGTGGGTTG TACTTCAAACTCCACGGGGTAAACAGGCCTCCCGCGATGAAGAAAGATGGCATACAGAAA AGGAAAAGAAAGCCAAAGAATAATCAAGCAGATATCAAAAAGAAAGGCCTGTCACCCATA TCAACAGGTGCGTCTAAGAGTTCCAATGAAGTCAGCAGCGGGAGCACAGCTCCAGCTTCT TCAAATTATTCGGGCATAGCTTCCTTCGCCACCGGAGTATCAGGAACGCCTTCAACACCA TCGTCTTCGTCTGCCGCAGTTACGTCTATTGTAGATTCTCCCAGCCACCAACATCACAAT CATCAGCAGTCATCTCCGTTGACCCCTTCTTCATCATCCTCTTCATCATCACCTTCTTCG TCGTTGTTGCAAAGACATATGCAGGGTCTTACCATGGGGGGCCATTTAGATCATCACCAT CACCACCACCACCATCAT

>TRINITY_DN78099_c0_g1_i1 ACTCTGAAGTGTATAGAAAAGGTAAATAAAATTCAAGCCTATTGTGAAGAGAATGAGAAT TATGGTTTTATATAACAGAGAGCAATGAATACAAAGGTAATATTCAGTGATTGTAACATA GTAAAGTTTTTATTGCAGCTTTATTAACACCTGCACAGAAATAACCCTGCTGTGGCTCAC TTTATATGAGGAAGTGCTGCATTTTTATTGGCCACAACACAGATATTCATGGCATATTTT TGGAATTTTTTTTTTCTGGATACATTTGAGTTCTAAATGAATACATATCTCACTTGCTGG ATGGACTTCGGTAACTGAGCATTGCTTTGCTGTGGTAAGTTACGAGACTCCGTAAAGCAC AGTTTGATGAACACAGGCTGAAAGAGAAGTTTTTCCAAACTTAGTACCTAACATTTTATT TGGGATTTCAAATGTCAAAATCTGTTATTTATGGCCAAGTGTGCATGTGAGGTATAAAAT GAGATTCTTTGCCATTTGAACAAGGATTGGTTTGTCCTGAAATTGTTATGGACGATACAG GAGGTGGGTATTCACTCTGTATGGCGCCTGCTGCAGACTGTGTTGTCATTGTAGTGATGG TATCTGATGAAGATGATGGGACTGGCACAGATGGTAGGTGGCATGTTGATGTGACTGCTG ATAAGGGAGATTCAACAGGACTCCTCACTTGTCTTCCTGACGGTGGTGAAGGAGTTGAGC TAGGTGGGGCACTGCCATCAAAACTTAGCTTTTTGTCCTTCTCGGATGAGCCTTGTACAT CTGGGACTTCGGAAGACACAGCTGCTTTGGGTGTTGCTGTTTTGGGCTTCCTTTTCCTGG TTTGAATTCCAGCTTTCCTCATAGCAGGCGGACGGTTCATATTGTGCAGTTTATAATAGA GACCACATGCGTTGCATACAGGTTCTCCGTGATTGTTTCTCCTCCATAGTGTGGTGGTTG TAGTTCCACAGTTCGAACAAGATAACCCCAGACGTCGTGTTGCTGGCAGCCTCCTCTGAG GTTTCGACAGTGGCCTGTTGACTCCGTTCATCCTGTGGTATAATCCACATGCATTGCACA AGTAGTGGCCGGTTCCATCTCTCCTCCACAGGGGGGTAGACACAGCTCCACAATTAACAC ATTCTCGGCCTTCGTTGAACTGAGAATAATCGTGAGAATCTGCTCTTCGATTTAGGTGTG CCATTGACGATTGCATTCCATTGCCGTTCTCGAAGCTTGACCAATGTCCGTTGTCCGTGA TGTATGAGTGATAAGGTGAGAATGCGTTGCTTCTTTGAAAATACGAGCCTCCGTCCCTAG TTTGTACCCCCATGCCTGGAGTTTGGAATCCAAACCTGGTTGTCGATAATGAGCTATGCG CAGACGGATGATACGTCTCGTGTACCTGAACGGCAGCAGTCCTATTTGGATCACCAGTTT GACCCCACGGGCTTGGGTTCGAACAAGCCATGGTTGCTCCTCCTTGATGCAGAGTGACCA TTGGCCTGGATGCTGGTAGGAATACGGGACTAGCGTGAATGTATCCGTCTTGATAAGATG GCATGTGGATAGAGGACATATTCTGGAACATGTTGGGTACATACATATATAGAGAAGAAG CAGTTATCTTTATCGGCCGAAACCGTCTCTTGGGAAATAAAATACAATTTTTCGTGAAAA CGCTGACTTAAATGCCAATGCGCAGTGTGCCGCTGGATAAACTTGAAAACAGTAACATAT TTATCTGTTTTATAAGATGGAGAGAAAGAGAGGTGCAGGTTGCAGAAGAAACATCACAAG CAGCTTCGTTTACCACACTCTGTAACTATGTCATAAAAGTCCTTGAAGTTCAACTTCCTT TTAAGTTTGATCATGTGCTTATTCTACAAAGTTATTTAGAATTCCAGTTTCTGTTACAGC AGAACACTTCTTGAAAGACTCAAACACTATTTAGTTGTTTCTTTACTGAAGGAAGGGAAT ACTAATCTTTTTCTTCCTGATGTGAGATCGGCTTGAAAGTAGATAAACTTGTCCTATCTC

TGTGGTCTAATTTTTTTATTTTAACTCTGTCACAGAATCACTTATAAAAGATCACAATAT TCTGTAGAATAATGGTATAAGATTATCTGTATGAGAAGAGGAAATATCTTCTTATGATAA AAAAAGAGTAACTCTCCTCTGGTAAAGAACCAGGAACTCTACATCAGTATGCAGATCTCA AGAATCTGAACCAGACTTTTTCTCATTCATGACTTTTTCTCAAGAATCTGAACCAATGAC TTTTTCTCATTCTAGACTTTTTCTCAAGAATCTGAACCAGACTTTTTCTCATTCATGACT TTTTCTCAAGAATCTGAAC

>TRINITY_DN80076_c8_g1_i1 GGTGAGGCGAGAGGACCTCGACCCTCGACCCCATCGCAGGTCCAAACTGTGCTTATTTAC GCCTAAAACCACGTGGTCAGGGTGTAGCTGTATCTCGCATAAGTGGCCCCAAACAGTCCC AAGTAACGAACGATTTCGGAAGGAAGGAAGCAAGCCTCCCCGAGGAAACACGGGAACTAA AATGCGATGGGATATGGTCAAGTCTCAATGGTCAGTCAAATAGCCACTGATGATTTTCCT ACTGTTTTAAACATCGGGATAGCGGAGCATGATACTTCGGTACGTTCTTGGTTTATTCTT TGATTGAAGCAGAAGTTATCAAAGCAAGGTTATCGTTAGACGCAAGAAAGTTGCCTGAAA GGGATAATGTTATCATAGTGGTGCCATGGCGAGGTCTCAGGCAGAAGTGACAAGCAGACA GAATTTCTACCATCACAGGAGAGTGCAAGAATTGGAGAAGGATGACAAAGATGTTAATGA ACACGAGGTAGAAACAGAACATAGCGTCCCAGCTGATCTTTCTCCTCACGTCTCAGAAAT CAGAGATCACGTGAACGAGCACATTTTACATCACCACCATTCTAATGAGCACCGGGATAC AGAACAGGATTCTTTACTCCAGGACTTCCACGGACAGGGTGAACAGCGGGTAACACATTT AGCTAACCTCGATGAACAGGAGCGACGTCATCATGACCATTTAGATGCAGATGCAAAAGA CTCTAGTAATACAAGTAGTATCCCATCTTATAATCCGGAGTCGGGAGAAACTCCCAATAT CCATCATCCCCCTCATCATCATCATCAAGAGCATCTCTTTTCGGCAGATGATAACCAACC AAATCACATTCATATCCAGCATGAGGACCAACATCACCATCATCATCACTCCACCCATCA CTCTCATCCTCATCAGAATATTTTGATGCATTCGTCGAATTTCCACAGTGCGCTCCATGA ACAACAGCAGCAGCATGACGAAGATTCAGCAATACCTAATCACATGAATGATGATAATCA TCATCTCTCCCATCTTGAAGACGTAGCAGGAGAGGGAGCTGAAACAGCCCAAATGCATCC TCATATTTCTCACCAGCCTCATCATCTTCACGCCTCTCATCATCACCTTCAGGATCAAGA CGATCCAACAAACTCTCACAACGCTTTACAGCATCACACCAGAGATATTCTTCACCATGC GGAGAAGGAACTACAAGCCCACATGGATTCACCGGGAGAAGATCCTCGTATAAGCCTTGA CCTCCCTGAGGTTCACGGATCATTCCCGCACATTCGAAGCCCCACCTCTCCAAGAGATTA CATCCAGGATCGTCTTGCTACATATACTTTGCCCTCGGACTACAGAACTTTAGCAAGTAT AAATTCTTCTGATCCTCAAACAATTGAATCTTCACCGCTGAGAAATATGCTGTTCCCCTC TATGGGTTATTTGTCATCGTCACCCTTAACTTCTCAAAGGGATATTGGTGTTGATTCAGC ATTACATACCGTTAGTACACCTTCTGTGACATATCATCATTTGCCAGTAGCATCGGAATC ACCTGTTAGTCAAAGTGTTGCTAGTTCTTTATACGTGCCAACGGTATCTACACCCTCTAA ACTGTTAGGCTTGTCATCTTACGGAAGAATTCACGAGCATCCAAGCTCAGGAGCCGGGCA CTCTCTAATGTGGAGTCAGCTGAGTGAAGATCTTTCCCCTAAAACACAAATGTCCCCTTC GTCATCATCAGTATTATCCAGAAATGCTACAGTCGGCCATTCTTCTTCATACGTCAGCCA AGATATTTCCTCATGGGGCTATGACCCGAGTTCAGCAAATATGACTGCTCTTCAGTTGTC AATGGCATCTTCGCACTCTCCACCAGATGGCGACATGTTTGGCAGCCTAGAGAGCCGCGA ATGTGTCAATTGTGGTGCTATTTCGACGCCATTATGGAGGCGGGACGGAACTGGTCACTA CCTATGTAATGCATGTGGGCTTTACCATAGGATGAATGGCATGAATAGGCCTGTAGTCAA GAATCAAAAGAGGCTGTCAGCTTCCCGAAGAATGGGTCTGTACTGTTCTAACTGCCAAAC AACGAATACTTCTTTATGGAGAAGGAACAGTCAAGGGGAACCAGTGTGCAATGCCTGTGG ACTGTACTTCAAGCTTCACAGGAAGACATGAAATACAGTTTAACAGGAAGACCAATGGAG ACAACAAGTTTGTTAAATTCCTCGCAGTCATATTTCGTGTATACCAATACCAGTACAGGA CAAAGGACTCTTCCAGTCTTCACGTCAATGAAAGACTTGCCGAAATCTTCTTCCACTATG CATATTGTGAGTCCACCTTTACTTCATGGCGGTGGCAACACTAGTAATCTTCCTTTCTTG

CTTCAGACAGGAAGTAACACTGTGGTTGGATCACCGTCTTCCCCCAAGCATGAGCAAGGC TCGGACTATCAGATTTCGTCAACTGGACACGCCACGTTACAGCAGGCGCATGTGCAAATT GCAGGAACAGAAGGAACTCCCAATGACTTTGACCAACAACATGATCTGCTGCCGTGATTG GTGAAAGATTCTGGACGGTCACCCGCTTTCTTGATGACATTTTCTACGCCTGTCAAAAAA CGAGGGGAAAAATGCCATGTCAGTTGTCGTGTCAAAATGTGTGCGTCGTTTTAAAAATGT ACATAAAATGAGTTAGAGGAGAATACGACGTGTAAGGGGATTAAGACTAAATCGTCTAAA ATGTATGTATTTTTTTGTTTTTCGTTCTGTTTTTTTTTTTTTTTTTTTTATGT

>ara.DN78359_c3_g1_i2 CGGGAGACGCCGTCGAACGCGAGCGGTCGTCTGCAACCGGAGTAACGCCAGACCCAGTTC AGGCGTGGCGAAAGCGCAAGCGCTCGTCCGATCGAAGCTTAGTATCTTCCAGTCGGCGGC ACAACGCTCAGCGGAGCAGGAACTGCTACCACACACCGCATCGAAGACCATTCCCGCGTC CGTGCGATCTTCACTGAGCTACCCACCCCCTCCTTTTTTTTTCCATTGTCGCGCTTTAAA AAAAAAAAGGAAATCAGTTGGACAATGGACATCTAGTGGTCTTTCGTTTCTCCTCTAGGT TAACAACAAACGGAATACTCAAATAAATCGTCACTTTCTGTGCGAAAGGTAAGCGATTTC GTAGCAATTTCTCTCGAGTACATATTCGAAAATTACAAAACTGAAAGAGACGAATATCTT AGAAGAAAAAAATACTTGTGACGAAGAGTTGGTGGATTTTTTTCATGCAAAAGAACAGCT CATTAATGCATTTATTTGCCATCATGGAAAGTGCACTGTGAAAAATTATATCATCTTTCT GAAAGTTCAAAAAGGGCTGAAACAATTTTTCCTGAAAAAAAAAAAAACACCTTTAATAAT TAAGCTTCTTTTTTTGCCAGTTAATGTGAACAGTATCTGCAAGATCTGTAATAAGTTGTT CATGGTAGACAATATACCTGGAAAAATACAAGAAAATTACAAATTTGAAAAAAAAAACCC TTCAATGTTGTAACATTACAATTAGAATTTTTCCTGTCATGCTGTAACTAAAACAGTGTT TTGAATTTTGTCTCGGAGCATGTGACTTAACTGACACTTTTCGGACAAAACTTGTTACTA ATTTAAAAAGTTTAAACTTCCTTAAAAGGTGAATTTGAAAATTTAAAAAGACCATGTGAT CTTGAAGTAGCAGCCAATAGTTGCTGCTGGTTGTAAGAAATCGAAAGAAATAGACTGAGC GACTTTCTGAGAGAGAGAGAGGAAAAAACAAAAAAAAGCACATCCCCTCCCCCTCTAAAT GCGTCGACGATTTCCTCTAAAACTGTTGGACGCATTTTGCTTCATGGCGTGACATCAGCT TGCTTAGACATCTTATGTTCTGCTATTTCTCACATTGCGTGACTAGAGTGAAAATGGTCT TGAGTTAAAGGATTTTGTCGCTGAACGCCCACCACGAATCAATGGACTCTGGCATGAGAG TTTCTACTTCTTGAAGATGTCATGTCCTCAATTTGGTTATAACGGGTCACCTTCTTCAGC GCAGGAAAAATAAAATTGAATTGAATTGAACCGAGAGATAGAACAGACTCTGGGCATTCA TTCGACACAGCTGCCAGAAATCCTGCACGAACATTTACGCCTCAGGAAGGTTTCGTGCCG TATACAAATGCATCTTCATCTGTACCCTGTTCATCTCTTTTGATGGCCGGGCAGTCGGCG ACAGGGCAGCAATTGCATGGTGGGTCCGGAAGTGGCACGGGCCAGCCGTGCTGTGAATCC GGTCGCCCCGTGCTGACGGACCCCCTCACGGGGCAGACGGTGTGCTCGTGCCAGTACGAC GCCCAATTGCTTAGCTACCAGCGGTTGGCTGCTTCGGGGTTGCCGCTCAATATGTACGGC ACGGCGGCTGCAGCAGCGGCCGCGGCGGCGTACGGCGGAGAGCAAGGCTTCCTACCGCTC GGAACTGAACAGTCAGCTTTCTACTCGCCATCGGCAAATGGCTTTGATCTAAAAGACAAT CTAGAGGCTTGGCGGAGTTTGCCATACGCAGCTTCTATGTATTATCCGTACGACTCTGCA GCACTAGCTGGTTATCCTTTTCCTAACGGGTACGGCGTCGACCTCAACGGTGCACGGAGG AAAAACGCCACGCGAGAGACGACGAGCACTTTGAAGGCGTGGCTGAATGAGCACCGGAAG AATCCTTACCCCACAAAGGGCGAGAAGATCATGTTGGCTATCATCACCAAGATGACCCTT ACCCAAGTGTCCACGTGGTTCGCCAATGCGAGGAGGCGGCTCAAAAAAGAGAACAAAATG ACCTGGTCTCCTCGTAATCGTTGCGACGACGACGATGCCGATCCCGACCCGGAGGACACG GAGAGACCGAAAGATGTCGACAACAAAGGCCGCGACAAAGATGGCGTTGATGCGGATCGG

### araucan

CAGAGGAGGGATGCCGGGGATAAGGAACCCGACACGAGGCAAGGAGGTTCCGATGATGGC AAAGTTGATTTTGTCGACGTTGACGACGATTGCAAGTCATGTGCATCAACAAGTACTGAT GGCCATATTATTAACTCTTCTGACAAAGAGAACTGCGTGTTAAGTGCTAGTGGTAAACCA AGAGTTCCCAGTCCTGTATCAGATGCCTCCCTCGCCTCAGACTCTGAGTCCTCTCAGGCT TCGGTACTAATGCCCGGTGTTGTAATTAGCAGTCAGCAGTCAACGGAACTTCCGGCCAAG CCACGGATCTGGTCTATAGTGGACACAGCAACCTCAAACACTTCTGCGCCGAGCACGTCA GTGAACTCTCCTCCCAGTGGTGGTGCTTCTAGACTAAGTCCCCTGCAGAGGGCAGCTTAT CTTAGCCCATATCCTAAACCTACATCGTGGTATGCTGGAACTTTAGGTAACCTAAGTGCA GTGACCAGTGGAGTAGGAGGAAGTTTCCCGTTGTCAGCAGCAGCCTTCACTTGTCCGCCA ATAGTATCTTCGATGACAGGTTCTGTTGCGTCCTCTGGTTCATCGTCCCTTGTGGAATCC CCTGCAGTGTCTTCGCTAAATAGACTAAGGACCGCAGCGGCGGCATTTCCGACAGCATCT TCCGGAGAGCTGTCGTTACACAAGGCCGCTGCTGTCGCAGCTGCAGCAGCAGCAGCTGCG GCGTCACCAACGGCTTCTCACACCCCTTCGGTATTCAACCTAAGCACTAAAGTTTCTGGA GGTCATTCCAGCTCGCATGTATCTTCAGGATCAGCGGCGGCCATAGCTGCAGTGGCTGGA AGTGGTAACACAGTGTCTTCCTCGAGCAGCGGAATATCGTCTTCGGCCAGCAGTCAAGGG AGAGTGACTTCGTGAAAGCGAATTTTTCTCGGTGATTCTAACTGTACAGTATAAAAAAAA AGATATGAAATAAAATTTCAGGAATGTAAATGTGTACCATATACTGTTGGGAATGGTTCA TTTGTTTTCTGATGATGAGACCAATGTGATTCGCTGAAGGATCTCTTTTACATGCCGTGG AAGATCTGCAAAGTTCCAAAGTGACCCATCGTTCAGTAAACCTCCTATGAATTTCGTACA GAGCAAACTACTAAGCACAAAACAAACACTTTAATCAGACTTCACAATTCCCTCAGCCTG TATCAGGAAAAACATTTTCTATAATTTCCTGCAGCATTTGTGCAACTACAATGAGTTGTA AATATCTTTCCTGAGAATGTACAGCGTCTAATTTTGCAACTCTCAGGAACGAATTACAAT TCGTCACTATGACTGCATTGAACAACTGAGAGAAAGAAAGAATTGCAGCGCCAATCAGAT TCGAACACTACTGTACACATGAATGGACTAGATTAGATAAGGAAAAGAAAGACCTTATGT AAAATTGTATTATAATATTGCAAAAGAGGAAATTATAAATTGTACTTCACAGTGTTTTGG CCCACCATCTGAGAAAGCCAAAATACTTAAAAGAAATCGATTAAACAGGCCAAATTGTTA TCGTTAAATTAATGCAGGATATAAGATATATCGTTATGATCGACATTATGCTTTTTCTAA ACCAAAGTTGTACATACTTTCTAATAAATATATGTGTACAAACGTTCAGCTTCGAGAACG TAAGCATGTTCTCCAAGAAGATTAAAAGAATAAATAAAAAGAAGTTATGACACGTAGAAC CTGAATATCATCATGGGTCATTGATTCCTAAACTCCTCCCCCAGCGGGCGCTGGGAATTA TTTATTATTTCTTGAACATTTAAGCAGGGTGTATAATATAGATCAGGCGGTATGATACCG CCCCTGATTTCACTGAGGAAACCCCGTGTCACCCTCATTTCCTGTGCTGCATATCCATCA AGTTATATGAACATGTGATTATGTCCAAATGTCAAAAAAAAGTAATGACAAACTGTTTAT GTCACAATAAAAGAGCAGTGTTTCGAAATAAGGAAAAAAAAA

### odd-skipped

>Acanthoscurria_odd-skipped_TRINITY_DN78272_c8_g1_i1 len=1979 AGAGAATTGTAAAAAGTGTCGTTCTCTAATTATTATTGAGTGATTTAAATGTGCTACAAT GCCACTTTATTGTAAGAACAGAGGGAAGTTATTGTGTGGGCATTCAAATAAGTACACTGT AGTAATAAAAGGTATAACTGTGAATTTTCAGGATCACCATATAAGATATACCCAGCAGAA AAAAATGTATTCCATTACAAGTCGTTTTATACGTAATAAATTTTATGTTCCATTCGTTTT CCTATGACATGTTGATTGTGGCATTTTGTATCATAAATACCTTATCTTTGGTATATACAT ATATATATTGAAGCACCAGTTTTCATGACCAAGTACCATGATCTCCATTTTTTTTCACAT ATATATAATTTCAAAATGTTAACTGTATAACCAATTAACTCTTGTTTGTCTTACATATAG AATTTAATTCAAATATACAGTAAAGTCCACCAGGGATTTTGATTGTCCTTTTTCCTTTAT ACATAAATATTCTAGAACTGGCATGTTCACAAACTACCTGCAGATGGGTGAACCAGATTT AGCGGATGATTATACGAAGTTACATCGACCACAGCGTCGTCAGAAGACTGATGAGGTAGT

CCATCACTTCGAAGGCTATGTGTGAGGCTGTGCCTCCGAAGATCGCAGTTCCGTCGGAAC ACCTTCCCACAGGAATCGCAGTTGTATGGTTTAATGTCCGTGTGAGTGAGGAGATGGGTT TTCAGGTTACTCCTTTGATTAAAGGTTCTGCCGCAGGTGGGACACTTATGAGGAGAGTCT TCCATGTGGAGTATCCTGTGTACAGCTAAAGTCCGAGATTGGCAGAAGCCTTTGCCACAC TCCGTGCATTTGAATGGTTTCTCTTTCGAATGAATGTACCTGTGATCTCTGAGATGGTCC TGTCTTCTGAATGCTTTGTGGCAGATGTCACATGTGTAAGGTCGTTCATCTGTGTGAGTC CTCTCATGGATAAGTAAGTTATAGGATTTGGTGAACCTCCTTTGACAATATTTGCAGATG AATTCTTTCTTTGGTCTCGATGACACCCTACCGCGGAAAAACCTCTTCTCGAACAACAAG GAACCATTGGCACTCCCAAGAGGTGATTTCCTAAATATGCCAGCAATGGCTGGATTGTAG AGGTGACCGAGAAAAAGAGGATGGGACGGGAAGATGGCTGGTTCTTGAAGCGGAGTAACT GTTTTTGGCACCTCGGAGGCTACCAGACTGTGAGGTGGGGATTTGACAGAGCTGGGATCT TCAGCATCCTCCTCATCAGATCTCGTGGCTGACTCTGCCAGATGAGCAAAATCGAATTTC CTTTTCTTCTTCTTTGGCGATGTGGGTGTTTTAGGCTGCGGAAGGACGACATCCTGTGGC GGCACGGGTCTCGGAGGCAATCCGGAATAGCTGACGTCGCCACAGTCGTACGGCGACGGA GGGAGATAGGCTGTGAAGGCCGATCCGCTGATGTTCAGGTGAAGAGGAATCGGGACGACG GGAGCCTGAGGAAGCATAGTCGTCGTTACCGCAGCAACAGCGGCCGTCGTCTCCCGTTGC GGTGGTGGCCAGGTAGTGTTGACAACCGTCGGACGAAACAAGGGTGGGGAGGGGCTGCTG TGACTACTCGTCGGCGTGAGAGGAGGCGTCGGCAGGAACCTTTGGTGGTGATGATGATGC TGATCTCTACGTTCATCTTCTAGCAGGAGCGTGATGCTATTTGCAGCAGCAGCAGAGTCC GGCATGGCGACTGCTGGGGTGCGGCGGGGTTACTCACACACTCTGGGGGCAAGCCAGCCC AGGCGGAATTCGGCAGTCTTCAGGAACAGCAGCAACAGGAGCCGGTTCCAAAGTGGTCCT GTGTCGCACAGGGCAGTCAGCTACTGACTGGTAAAGCGAGCAGCAAGAGAGCAGCGGTCC GGGACCGCGGAGGGGGAAGAATTGCTCTGCTAGAACCCGGATGTTGGCGTGGCTCAGCG

### Distalless

>Dll_DN74055_c5_g1_i4 AGCCTATCCTGGTAGTTTCAAGTACACGTTCGTCGACGTCCGACACGCGGGTGTGCAGAG CAGGGCAAAGCAATCTTCCTTCTTTTTCTCTCAGTCGGGGTCCCGCTAAGCCGCCACCAC GTTGAATTTTACGGAGTTCTCGGCCCCCTACCCGAGTATGGCGGGTACTGCGGATGGCCT TGATCAGGATGTGACCAACAAGTCTGCATTCATGGAGATACAACAGCAGGGTCTGGCCGG TATGCCTCATCATGCCCAAGGTTACCCTATCCGTTCGACCTACTCAAGCCAGCCGACGCA GCACGAGGCTGTGTTCGCCACGTCCCAACACCCCAGGCCGCTGGGGGCTTATCCGTTCCC TATGAACAACTCACCTTTGAATACAAGTTTGCACAATTCCTACACACACCCAAGTCATCC ATATCTCAGCACATATCCAGCAAATGTTCCTGGTTGTCCACCGTGTCCTTCACCTCCAAG AGACGATAAGTCACAGTTGGAGGAAACACTCCGAGTGAATGGTAAAGGTAAAAAGATGAG AAAACCGCGCACAATCTACTCCAGTCTACAGCTACAGCAACTGAACAGGAGGTTTCAGCG GACACAGTACCTTGCCTTACCGGAGCGGGCTGAACTCGCAGCGTCCCTGGGACTTACGCA GACACAGGTGAAGATATGGTTCCAAAATAGGAGGTCCAAGTACAAGAAGATGCTCAAAGC TCAACAGCAGCAACAGCAGCAAAACCCCCAAGCACCAGGACCGAACCAGACGCCCGGAGC TACGCCGAATCTACAGCCGCCCAACCCAGCGTCAACGCCGCAGACCCCGCCTGAATCCCA CGACGCCCACACACCTCCCTTAGCACCGCTCATGAATCCACAGCAGCAGCAGCCGCCGCC ACCGCAGCAGCAGCAACCTCCGCCGCAGGGACCCGGTGTCATGCCGACTTCGGCCGTCAG TCCTCCTGCTATGAGTCCTCCCATCTCATCCTGGGACATGGCATCTTCCAAGGCAGCAGC AATGAACACTTATATGCCTCAGTACTCTTGGTACCATCAGACGGACCCCTCAATGAGCCA ACAAATTCTCACTTAATCCATTTTAAAACAAATAAGAATCATTTTTCCTGAGATGACGAA AATGAAGTGTGAATGAATATATTCGTGTGTGTTCATACAAAGGCTTCCCCCCCACTTCAG ATGGGTGGGTTGATGCATCACACGTTGGACATCGGCTGCCAACATCCCAGCCTAGTATCA TCACAGTATTATGTGACGGTGCGGTGTCATGAAGTGCCGAAGAAGAAGAAGGGCAGCCTG

TAAACAGCAACCCGTGCTCAGTCGGACGGCAGGATCGTGGACCGAAACATCCAAACGCCA GCCAGTCAGCCAGCCAGTCAGATAAAGAAACCAGTGTCACCCATTTCACAACCCTTCCTA ACAATCGTCGCAAACGACCTACTCCCCAGGCCCTTTTCTCCCCGAGTCCCATTCACAGAG GGACTTTACATCAACTAATATCAGCCATTTTAAAAATTCATCAGCTTCATTTGTGGAACT TCTTTGGCCAGTACTATATATACTTCTATGCCACAGTTCAACAGCCAAATGTAAACTTGA GAGGTAACAAGTCTTTCTTCGTTGTGTCCAAAGGGAACTGCATCCAACTCAAGAGAATGT CAGCAAACCGCACAGGATTGATGATCCCTGCTACACATCTAGTGAGATATTTCCAAAAAC ATCATGCCAGAATTATTAATTTATCTCGTACTAAGCACTGTCTTGGATATGTAATATCTA CTTGTTGTAAAGAAGATGGCAGTGTAGTTTTTGTACAAATGCTTCTTCGAGGTTTAGCTA TTTAAAATAAAAAAAAACAAAAACTCCATCTCTCAAGAAATAAGAAGTCTGCGACAGAAA TTGTAATGTATATTTAGTACAAATGATATCAATCATTCTTTTTCTTCATGTAAACAATAC AAAAATTGTTACTGAAACCGATTATATTATGCTAATTGTGAATGAATCTGTATCTATTTG TAAAATATTTTTATGGAAATTATTATAAGGGACAAGAAACTATATTGAACCTGATAAATA AGCGTACATATTATGCCTGCCTCAAAATTACTTCTTTTTCATGAATTAAGCCGCGAGATT TAATTTGTATAAAAGTCCACAAAATTGCGGCCGCTTGACATGCTCATGATAAATGCCACC ATTGCTGAGAAATGAATTCCAGTTACAATCGGAATATTCAATGATATTTTTACCCTCCCT TTCAATCGGAATTATTTCCATAACGCTTGGCCTCATGGATGACAGTTTCATCAATAAATG ATCAGGCACTGTTACAGATATCGCCAATCTTGCATCATACTAAAAAAAAAAAAACTAATA CAAGTTCATATTCATTGCAACTCGTTCAACCCAGAAGAAAACAGCAGCACAGACGAATGA TTTAGTTTTAAATATGCCACAAATCAGGTAACAGATTCAAAATTACACGGAAGCAAGAGT TGTACCAAGTTTTCTTACCTGAAAAGATTACGAATGGTAAATAAAATTCTTCTGTGTAAA TAAGTGGTCATATGTTCGAGCTAAGTCATATCTATAAGACAAATTAAATAGCATAAATAC ACTGAATTATGAATCATCCCGCATTAGTACACAAGAGCAGTTTATATTTTAATTACCGTA AGAAATATACAGAAGAGTGGTGGCAGAACTGCGGAATAACTTTAACACAATTAATTAATA AAAATAAAAAAAAATCATACAAGATATTTTTCTTACAACGTTAGTTACTGGTAAAAGTAA GCATGGTAATTATGCTTAACACCAAATAATACAGTAGACAACTATTTCGCCTAATATCAA

---

**Fig. S1.**
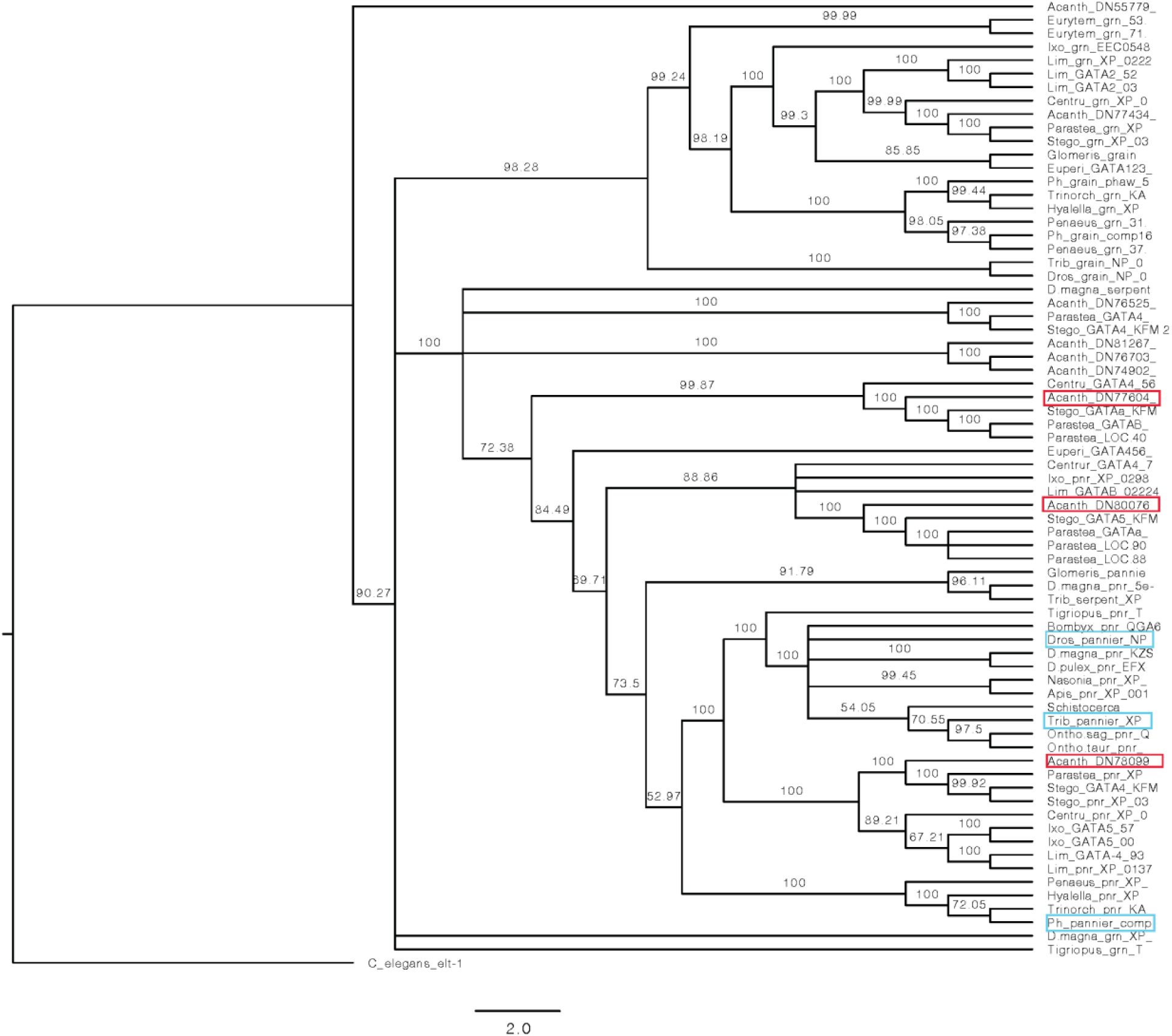
Three orthologs of *pnr* were identified in *Acanthoscurria* with closest homology to *Drosophila*, *Tribolium*, and *Parhyale pnr* (Fig. S1-2). However, only one of these was expressed at the stages examined, Acanth_DN77604, and was presumed to be *pnr*. Consensus tree generated using Mafft, which gave similar topology to Clustal consensus tree.

**Fig. S2.**
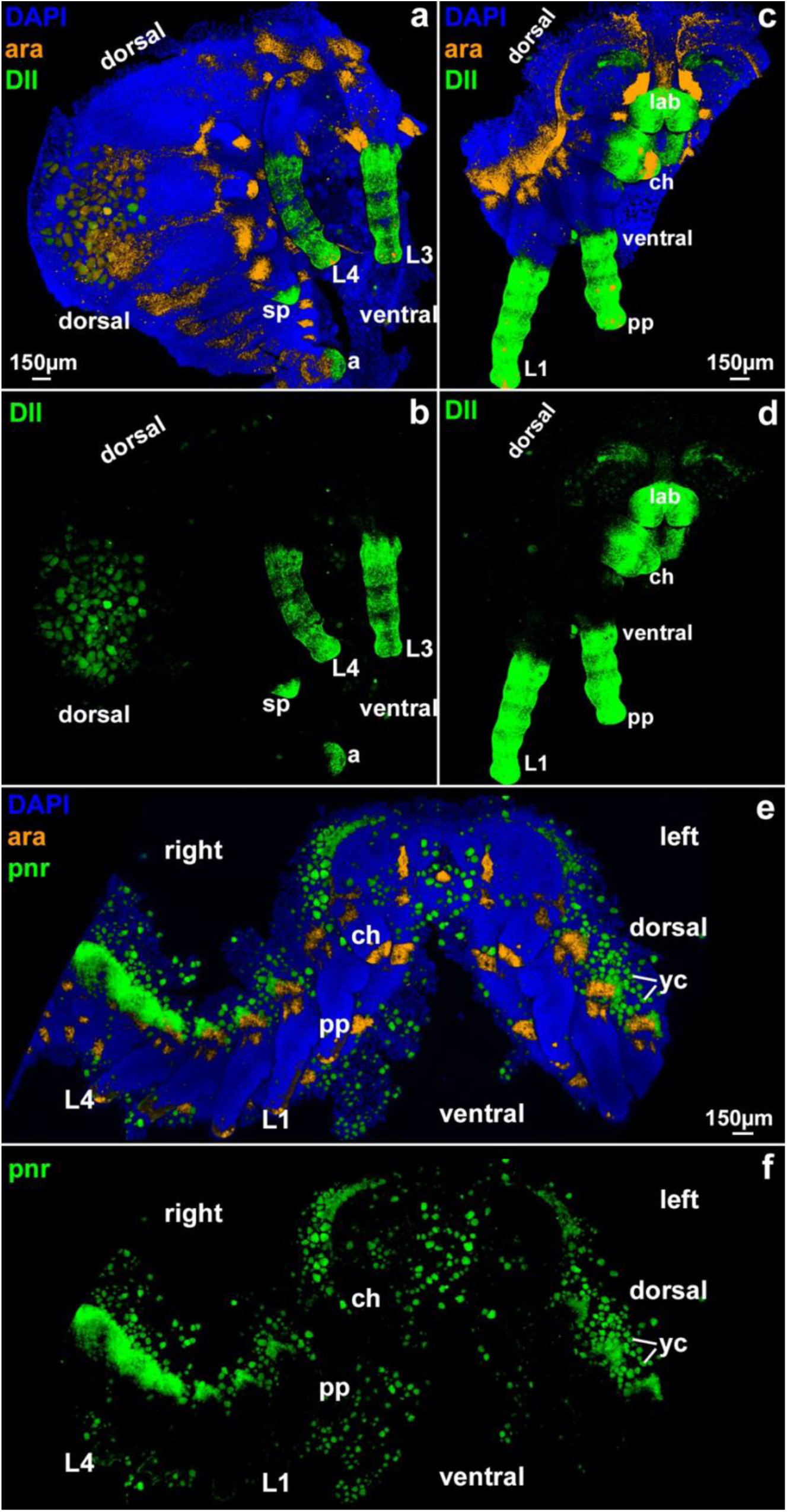
In situ HCR of separate *pannier* and *Distalless* channels showing no overlap of expression in legs.

**Fig. S3.**
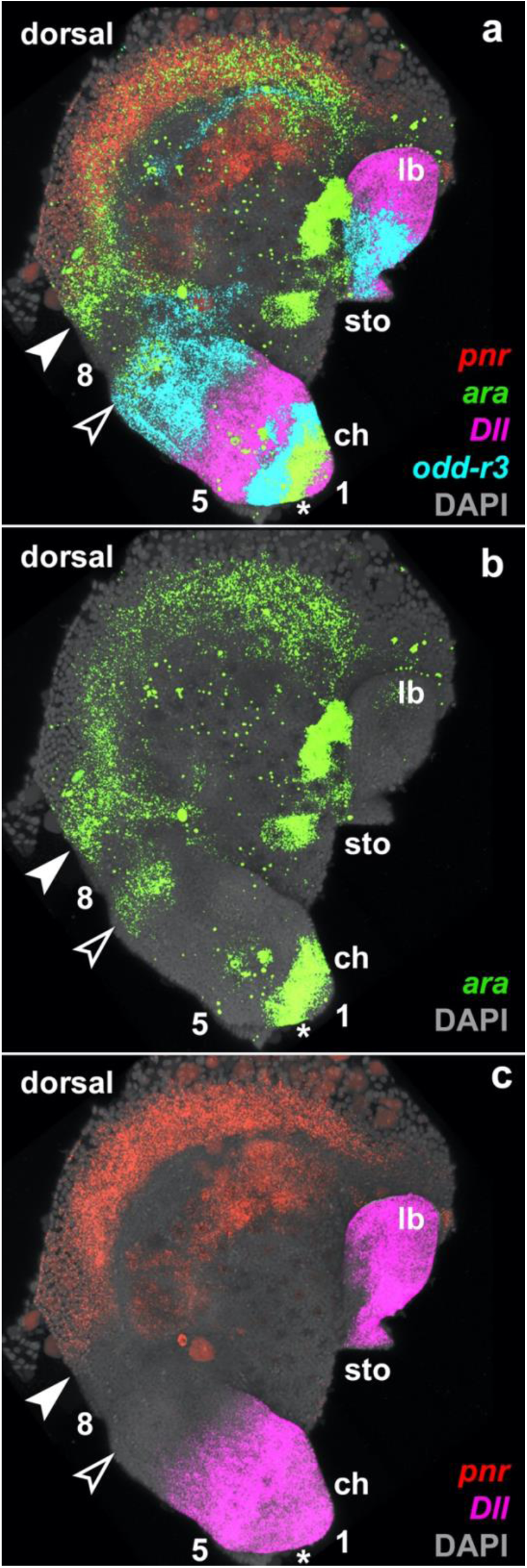
Leg segments in *Acanthoscurria* chelicerae. The expression of *pannier, araucan, odd-skipped-related-3,* and *Distalless* suggests the presence of 3 leg segments in the chelicerae, segments 1, 5, and 8.

**Fig. S4.**
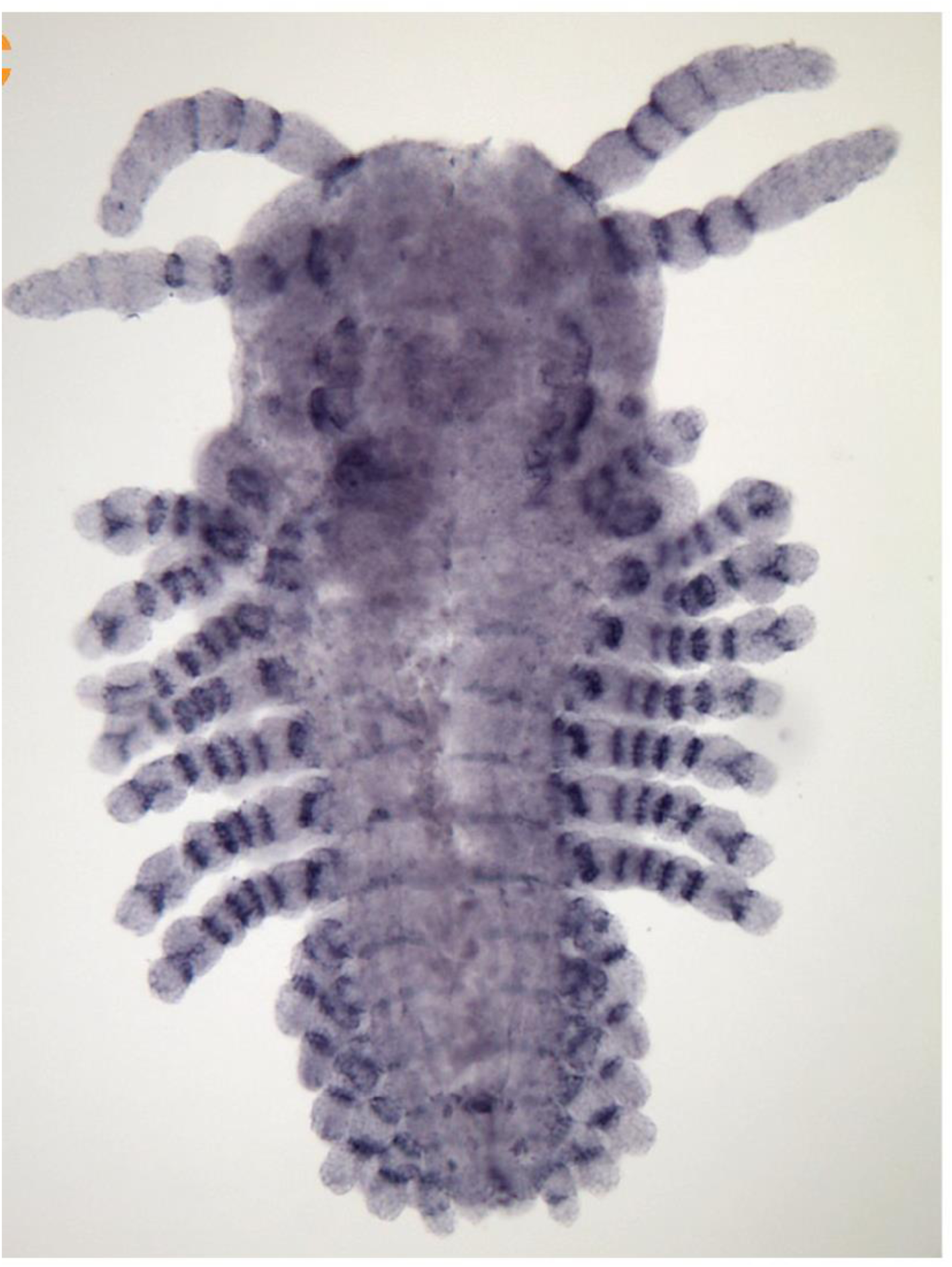
Expression of *drm* (*odd-skipped 2*) in *Parhyale* St 22 embryo reproduced from Kira O’Day’s thesis^67^. In the full-length thoracic legs, seven stripes are visible (six strong stripes and one proximal, fainter stripe that is most visible in the legs on the right), which separate 8 leg segments.

